# Bark from avocado trees of different geographic locations have consistent microbial communities

**DOI:** 10.1101/2020.08.21.261396

**Authors:** Eneas Aguirre-von-Wobeser, Alexandro Alonso-Sánchez, Alfonso Méndez-Bravo, Luis Alberto Villanueva Espino, Frédérique Reverchon

**Author notes:** Correspondance: E. Aguirre-von-Wobeser, CONACYT – Centro de Investigación y Desarrollo en Agrobiotecnología Alimentaria, Centro de Investigación y Desarrollo, A.C., Blvd. Sta. Catarina s/n, Col. Santiago Tlapacoya, 42110, San Agustín Tlaxiaca, Hidalgo, Mexico, 52-5525-24-8626.

## Abstract

Bark is a permanent surface for microbial colonization at the interface of trees and the surrounding air. However, little is known about the microbial communities harbored on these tissues. Studies on bark microbial ecology show a dominance of bacteria from a few phyla. Bark microbial communities of avocado (*Persea americana*) could have implications for tree health, as a first barrier for defense against certain pests and diseases in this economically important species. We used shotgun metagenomic sequencing to analyze the bark microbial communities of avocado trees from two orchards, and compared one of them to rhizospheric soil. Our results show that the microbial communities of avocado bark have a well-defined taxonomic structure, with consistent patterns of abundance of bacteria, fungi and archaea, even in trees from two different locations. Bacteria in avocado bark were dominated by Proteobacteria (particularly Alphaproteobacteria), Actinobacteria and Bacteroidetes, consistently with bark communities in other trees. Fungal members were dominated by Ascomycota and Basidiomycota, while most Archaea in bark were Euryarchaeota. We can conclude that avocado bark is a well-defined environment, providing niches for specific taxonomic groups. The present in-depth characterization of bark microbial communities can form a basis for their future manipulation for agronomical purposes.

## Introduction

Plants and microorganisms form complex systems, where interactions can result in beneficial, neutral or detrimental effects on the plants’ health and productivity (Mendes *et al.* 2013; Hassani *et al.* 2018). The most diverse and abundant microbial communities associated with plants are located at their roots, and have thus been the main focus of plant microbial ecology (Berendsen *et al.* 2012; Jacoby *et al.* 2017; Aguirre-von-Wobeser *et al.* 2018; Delgado-Baquerizo *et al.* 2018). Nevertheless, evidence that microorganisms living on the aerial parts of plants are important for their health is accumulating (Stone *et al.* 2018). While the rhizosphere environment favors the development of large and diverse populations of microorganisms, the aerial parts are a harsher environment (Vorholt, 2012). Exposure to ultraviolet radiation, desiccation, and scarcity of nutrients are among the challenges for microbes in the surfaces of leaves and stems (Leff *et al.* 2015, Stone *et al.* 2018). Furthermore, plant tissues such as leaves, and especially bark, commonly have protective chemicals, some of which show antimicrobial activities (Ogundare and Oladejo, 2014). Therefore, microorganisms capable of growing on these surfaces are likely to be tolerant or resistant to several stresses.

The bark environment, referred to as caulosphere (Garner, 1967; Nelson, 2018) or dermosphere (Lambais *et al.* 2014), is relatively stable, as compared with leaves, flowers and fruits, which are constantly renovated or appear only seasonally (Vitulo *et al.* 2019). Accordingly, within individual trees, the trunk bark has more diverse communities than newer branches or leaves (Leff *et al.* 2015). Bark microbial communities have been observed to host plant beneficial, neutral and pathogenic fungi and bacteria, and could act as a reservoir of these microorganisms (Buck *et al.* 1998; Martins *et al.* 2013; Arrigoni *et al.* 2018; Arrigoni *et al.* 2020). Known factors which impact the composition of bark microbial communities in fruit trees are the age of the plant, with older plants having in general more diverse communities, the plant genotype (species and cultivars) and the geographical location of the orchards (Arrigoni *et al.* 2018; Arrigoni *et al.* 2020).

The barks of trees are considered the first line of defense against many pests and pathogens, with constitutive defenses and inducible responses to attacks (Franceschi *et al.* 2005). Nevertheless, very few studies exist on the microbial ecology of barks of wild (Lambais *et al.* 2014), ornamental (Leff *et al.* 2015) and crop trees (Arrigoni *et al.* 2018; Arrigoni *et al.* 2020). Avocado (*Persea americana* Mill.), a native crop tree from Mexico and Central America, is an economically important commodity in Mexico, as the country has a share of approximately 35% of the worlds production (FAO, 2016; Rendón-Anaya *et al.* 2019). However, avocado production has been hindered by the incidence of several fast-spreading diseases, mostly caused by pathogenic fungi and oomycetes (Méndez-Bravo *et al.* 2018; Guevara-Avendaño *et al.* 2020). Surprisingly, avocado tree bark as a microbial environment has received little attention in the scientific literature, despite being the first point of contact for different pests and diseases. For example, stem and branch cankers can invade through injuries on stems surfaces (Guarnaccia *et al.* 2016). Most importantly, tree-killing fungal diseases are inoculated by scolytid beetles boring holes into avocado trunks and stems (Harrington *et al.* 2008; Eskalen *et al.* 2013; O’Donnell *et al.* 2016). Invasive scolytid-fungi associations, *Xyleborus glabratus/Raffaelea lauricola* and *Euwallacea* spp. nr. *fornicatus/Fusarium euwallaceae* and *F. kuroshium,* have caused extensive damages to the avocado industry in the United States in the last two decades (Ploetz *et al.* 2012; Carrillo *et al.* 2015; Na *et al.* 2018), and are spreading to other regions (García-Ávila *et al.* 2016: Lira-Noriega *et al.* 2018; van den Berg *et al.* 2019). As these beetles first land on the trunk surfaces, the use of microbes that antagonize those fungal pathogens on the bark has been proposed (Dunlap *et al.* 2015; Castrejón-Antonio *et al.* 2020). In this work, we provide a first approach to avocado bark microbial ecology, by using shotgun metagenomics sequencing, in combination with phylogenetic analysis, to determine the microbial community structure – including bacteria, archaea, and fungi – of Hass avocado trees bark. We show that avocado bark microbial communities are consistently dominated by particular taxonomic groups of bacteria, archaea and fungi, even in trees sampled in different times of the year and geographic locations. Thus, our results show that the bark of trees is a well-defined microbial environment, providing niches for particular microorganisms. Knowledge on the structure of bark microbial communities of avocado is a necessary step in efforts for future managing of these communities for the benefit of avocado producers.

## Methods

### Study sites

Two avocado orchards in different regions of Mexico were included as sampling sites for this work. The first one was near the city of Morelia, in the Michoacan State, where most of the avocado production in the country takes place. The second one was near Malinalco, in the Estado de Mexico State, in an area where small avocado orchards are common.

The Morelia orchard produces avocado for commercialization, and is managed with intensive agronomic practices. It is located in the rural locality of Las Palomas, at 19° 31’ 55.956’’ N, 101° 10’ 48.0216’’ W, with an altitude of 2370 m above sea level. The orchard, measuring 7 ha, consists exclusively of avocado trees from the Hass variety, grafted on rootstocks from local varieties. As is common in the region, the trees flower three times during the year, allowing frequent harvests. Large amounts of agrochemicals are used, including cupric oxide (Hidromet Flo, Tacsa, Mexico) as a fungicide, Astro 34% Permethrin as insecticide (FMC, USA), as well as the foliar fertilizers Nutri Wunder 12-62-00 (AgroScience, Mexico), Syntek immunopotentializer foliar fertilizer (AgroScience, Mexico), CaB fertilizer (Dragon, Mexico), MAXI-Grow biostimulant (Agrozar, Spain) and Fixed-Gro (Agroscience, Mexico) as adherent. For soil fertilization, organic pig and cow manure is applied once a year, at the beginning of the rainy season (early summer). Maintenance pruning is performed every two months during the dry months, and every month during the rainy season. The soil of the region where the Morelia orchard is located is classified as Humic Andolsol (INEGI, 1979), which originates from volcanic ashes. Climatic data for the closest meteorological station where retrieved from the Global Weather Data for SWAT website (https://globalweather.tamu.edu/request/view/28397), for the period from 1979 to 2013. The local climate shows a dry season from late autumn to spring, and a rainy season in summer (Fig. S1A), which is characteristic of central Mexico. Yearly precipitation has an average of 887 mm (standard deviation, 160 mm). Maximum daily temperatures increase during the spring, but decline at the onset of the rainy season (Fig. S1B). Minimum daily temperatures are lowest in the winter, at about 5°C, and increase in the summer to around 13°C.

The Malinalco orchard, located at 18° 59’ 22.2828’’ N, 99° 29’ 40.4124’’ W, is composed of old trees (planted in 1985), which were abandoned 18 years ago, and recently managed again (since October, 2019) for commercial exploitation, when the trees were severely pruned in order to obtain young branches for fruit production. Avocado trees in this orchard produce the Hass variety, and are grafted on rootstocks from local landraces. The orchard is managed with organic practices, and no agrochemicals are used for pest control or fertilization; only manure is added to the soil. Organic products added to the soil prior the sampling date were *Trichoderma harzianum* (Natucontrol, Biokrone, Mexico), a preparation of mycorrhizae containing *Glomus geosporum, G. fasciculatum, G. constrictum, G. tortuosum* and *G. intraradices* (Glumix, Biokrone, Mexico), diatomaceous earth and zinc oxide. The orchard has 380 avocado trees in 3.5 ha, of which presently 150 are productive. Besides avocado, other fruit trees are sparingly present, including guava, apple and citrus trees. The soil of the area is Chromic Cambisol (INEGI, 1976), which stems from volcanic ashes. Pluvial precipitation and temperature patterns were similar to the ones found in Morelia (Fig. S1C and S1D), with precipitation being slightly lower and more variable from year to year (mean 760 mm, standard deviation 357).

### Bark and rhizospheric soil sampling

Bark and rhizospheric soil samples were collected at the Morelia orchard, on 4 September 2019, and at the Malinalco orchard, Estado de México, Mexico on 9 April 2020. Bark sections of approximately 5 × 5 cm were collected with a sterile knife at approximately 1 m above the ground, and approximately 2 g of soil were collected adjacent to the roots, at a depth of 10-15 cm. Six trees located in the same area within the orchard, approximately 15-20 m apart, were sampled at each orchard. Samples were kept frozen for transport to the laboratory and were stored at −80°C until DNA extraction. Samples for sequencing were chosen according to the quality and quantity of the extracted DNA. Rhizospheric soil DNA from the Morelia orchard did not have sufficient quality for shotgun metagenomic analysis, and was not included in library preparation for this study. The sequenced library consisted of four bark samples and four soil samples from Malinalco and four bark samples from Morelia. However, not all samples yielded sequence reads for analysis (See results section for further details). All extracted DNA was stored for future analyses.

### DNA extraction library preparation and sequencing

DNA was extracted from bark and rhizospheric soil samples using a DNeasy PowerSoil Kit (QIAGEN). For bark DNA extraction, the periderm (outermost layer) was carefully removed using a sterile razor blade, and inserted directly into kit vials, filling about 3/4 of the volume. For soil samples, 0.25 g were used as starting material, as per the manufacturer’s instructions. Extracted DNA was used for library preparation with a Nextera DNA Flex Library Prep Kit (Illumina, USA), adding indexes from a Nextera CD Indexes Kit (Illumina, USA). Shotgun sequencing was performed on a NextSeq 550 instrument (Illumina, USA) using a high-output kit, which generates sequences up to 150 bases in the forward and reverse directions.

### Bioinformatic analysis

Bioinformatic analysis was conducted mainly in KBase (Arkin *et al.* 2018). Raw reads were preprocessed with Trimmomatic v0.36, to remove any remaining Nextera index and adapter sequences, and filter low quality sequences with a sliding window (Bolger *et al.* 2014.). Host contamination was assessed by aligning the reads to the avocado genome (Rendón-Anaya *et al.* 2019) using Bowtie v1.2.2 (Langmead *et al.* 2009). The taxonomic composition of the microbial communities was analyzed with Kaiju v1.7.2 (Menzel *et al.* 2016) using the nr database from NCBI, with random sampling of 10% of the reads. Taxonomic composition visualizations were created with Krona (Ondov *et al.* 2011).

Alpha and beta diversity analysis was performed based on Kaiju-generated taxonomies using R (https://www.r-project.org/), with the package vegan (Oksanen *et al.* 2019), considering the prokaryotic members of the microbial community. Alpha diversity was calculated as the Shannon index, and genus-level richness was represented as the number of distinct genera observed, by means of rarefaction curves. Beta diversity was estimated with the Bray-Curtis dissimilarity, and Principal Coordinate Analysis (PCoA) was used for dimensionality reduction and visualization.

### Sequence availability

Raw sequences were deposited at the Short Read Archive, under BioProject PRJNA656796, with accession number SUB7881803. Raw data are also accessible at KBase at https://kbase.us/n/69195/32/.

## Results

### Sequencing

Bark and rhizospheric soil samples were collected from two Hass avocado (*Persea americana*) orchards, and whole microbial community DNA was extracted and sequenced. We obtained a total of 852 million sequence reads from nine samples (Table 1). Reads from the remaining three samples (one from bark from Malinalco and two from bark from Morelia) could not be identified, presumably because of incorrect reading of the barcodes, or low DNA content in the library. Of the obtained reads, less than 2.5% corresponded to the host plant, *Persea americana.* Due to the low abundance of avocado reads, no pre-filtering of these was necessary for the analysis presented here.

**Table 1.** Metagonomic sequence reads analyzed

| Sample | Orchard | Material | Number of reads | Totals | Host Contamination (%) |
| --- | --- | --- | --- | --- | --- |
| A1 | Malinalco | Bark | 89,422,774 |  | 0.06 |
| A4 | Malinalco | Bark | 133,710,702 |  | 2.18 |
| A5 | Malinalco | Bark | 88,331,034 | 311,464,510 | 2.21 |
| SA1 | Malinalco | Rhizospheric soil | 93,743,192 |  | 0.86 |
| SA2 | Malinalco | Rhizospheric soil | 24,336,266 |  | 1.52 |
| SA4 | Malinalco | Rhizospheric soil | 62,453,588 |  | 0.01 |
| SA5 | Malinalco | Rhizospheric soil | 190,127,986 | 370,661,032 | 0.01 |
| M1 | Morelia | Bark | 99,693,356 |  | 0.01 |
| M2 | Morelia | Bark | 70,359,818 | 170,053,174 | 0.02 |
| Total |  |  |  | 852,178,716 |  |

### Alpha diversity and genus-level richness

Alpha diversity was calculated at the genus level for all samples for the identified prokaryotes. We excluded eukaryotes from this analysis because many of them are multicellular and/or polyploid, which affects the direct comparison between these groups. As genus-level taxonomic assignations were performed using a reference database, only named genera were considered for this analysis. As expected, the rhizospheric soil samples had the highest diversity levels and genus richness, with about 800 genera with abundances above 0.01% (Fig. 1A). The bark samples from the Malinalco and Morelia orchards had similar richness, with approximately 600 genera, but the Shannon diversity index was lower for the Morelia samples.

**Figure 1.**
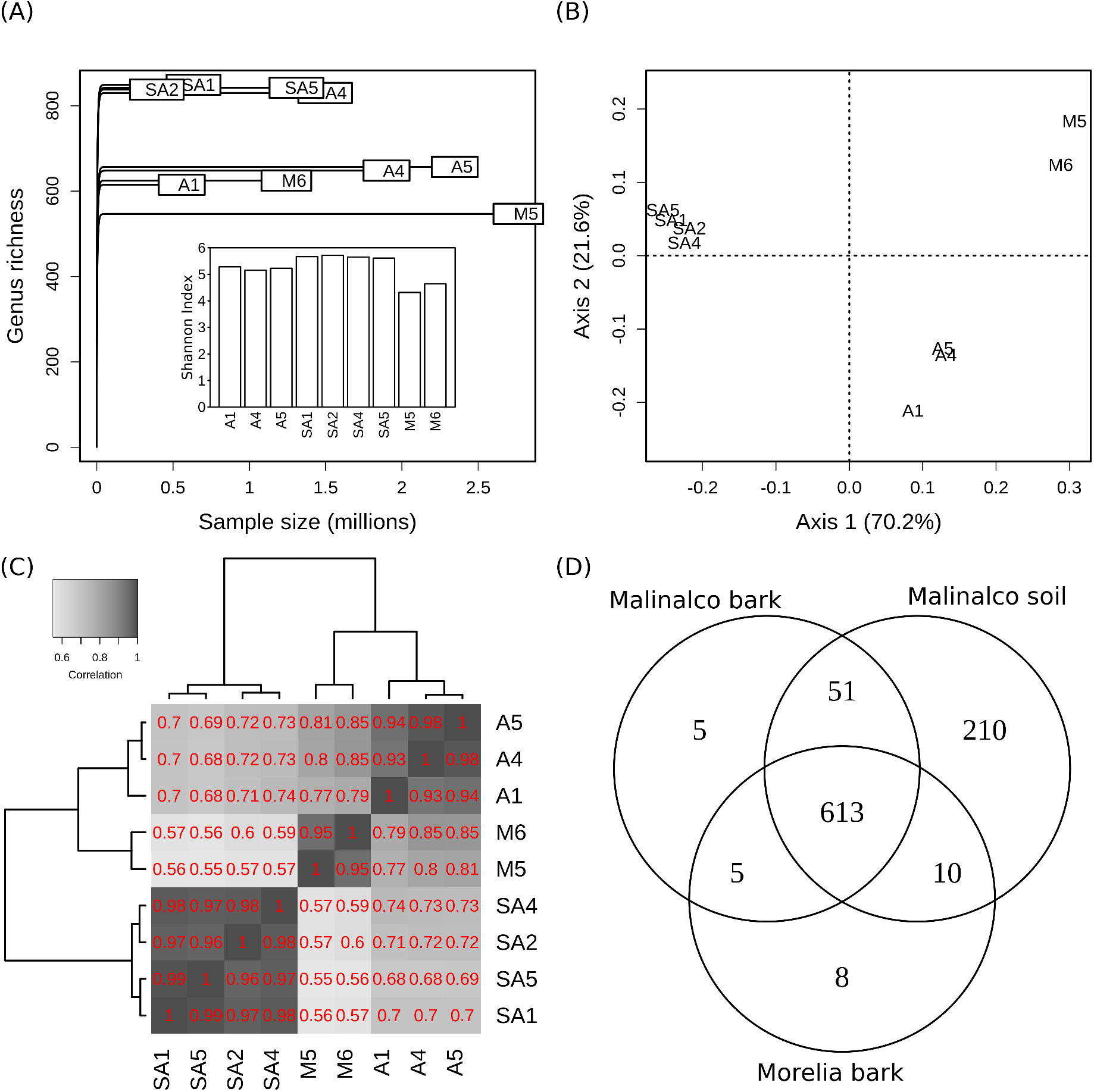
Diversity of avocado-associated bark and soil microbial communities at the genus level. Genera with an abundance of more than 0.01% were considered. A) Richness rarefaction curves and Shannon diversity index (Insert). B) Bray-Curtis dissimilarity Principal Coordinate Analysis. C) Correlation between abundances of genera in sequenced samples, ordered by hierarchical clustering of the correlation matrix. Relative abundances were square-root transformed. D) Venn-diagram of prokaryotic genera observed in bark from the Morelia orchard and bark and soil from the Malinalco orchard.

### Beta diversity

Based on relative abundances at the genus-level, the Bray-Curtis distance between samples was able to produce a clear separation of samples according to their origin, as shown by PCoA analysis (Fig. 1B). The first and second PCoA axes accounted for more than 91% of the variability, which implies that tree compartment (soil *vs.* bark) and environmental conditions (two different sampling locations and times of the year) are major determinants of the microbial communities in avocado-related samples. The abundances of prokaryotic genera in samples within each group were highly correlated with each other, with Pearson correlation coefficients above 0.9 for square-root transformed data (Fig. 1C and S2). Moreover, high correlations of around 0.8 were observed between the abundance of genera from the Malinalco and Morelia bark samples. Within the Malinalco orchard, the correlations between rhizospheric soil and bark samples were also large (around 0.7). Of the prokaryotic genera observed at a relative abundance greater than 0.01, the majority (613) were found in all three sample groups (Fig. 1D). A sizable proportion (210) were observed in rhizospheric soil exclusively, while the rhizospheric soil shared 15 and 10 prokaryotic genera with the Malinalco and Morelia bark samples, respectively. Very few genera (only 18) were found exclusively on bark samples, with 5 of them shared in both orchards.

### Community structure

The microbial communities of avocado tree bark were dominated by Actinobacteria, Proteobacteria (in particular, Alphaproteobacteria) and Bacteroidetes in both orchards (Fig. 2, S3-S10). Cyanobacteria were very abundant in barks from the Malinalco orchard, but were much less dominant in samples from Morelia. Other represented bacterial phyla in bark from both sites were Acidobacteria, Planctomycetes, Chloroflexi, Verrucomicrobia, Gemmatimonadetes and Firmicutes. Similar to the bark samples, Proteobacteria and Actinobacteria were also among the most abundant phyla in rhizospheric soil. However, Acidobacteria was the third most dominant phylum in rhizospheric soil, this being the largest distinction between the rhizospheric soil and bark environment in terms of bacterial phyla. The most abundant bacterial genera found in avocado trees bark from both sites were *Sphingomonas* (and closely related *Sphingobium*) and *Methylobacterium* (and closely related *Methylorubrum),* both belonging to the phylum Alphaproteobacteria (Fig. 3A). These genera reached abundances of up to 15% of all bacteria in the bark community. Other abundant Alphaproteobacteria were the rhizobiales *Bradyrhizobium* and *Bosea,* as well as *Roseomonas.*

**Figure 2.**
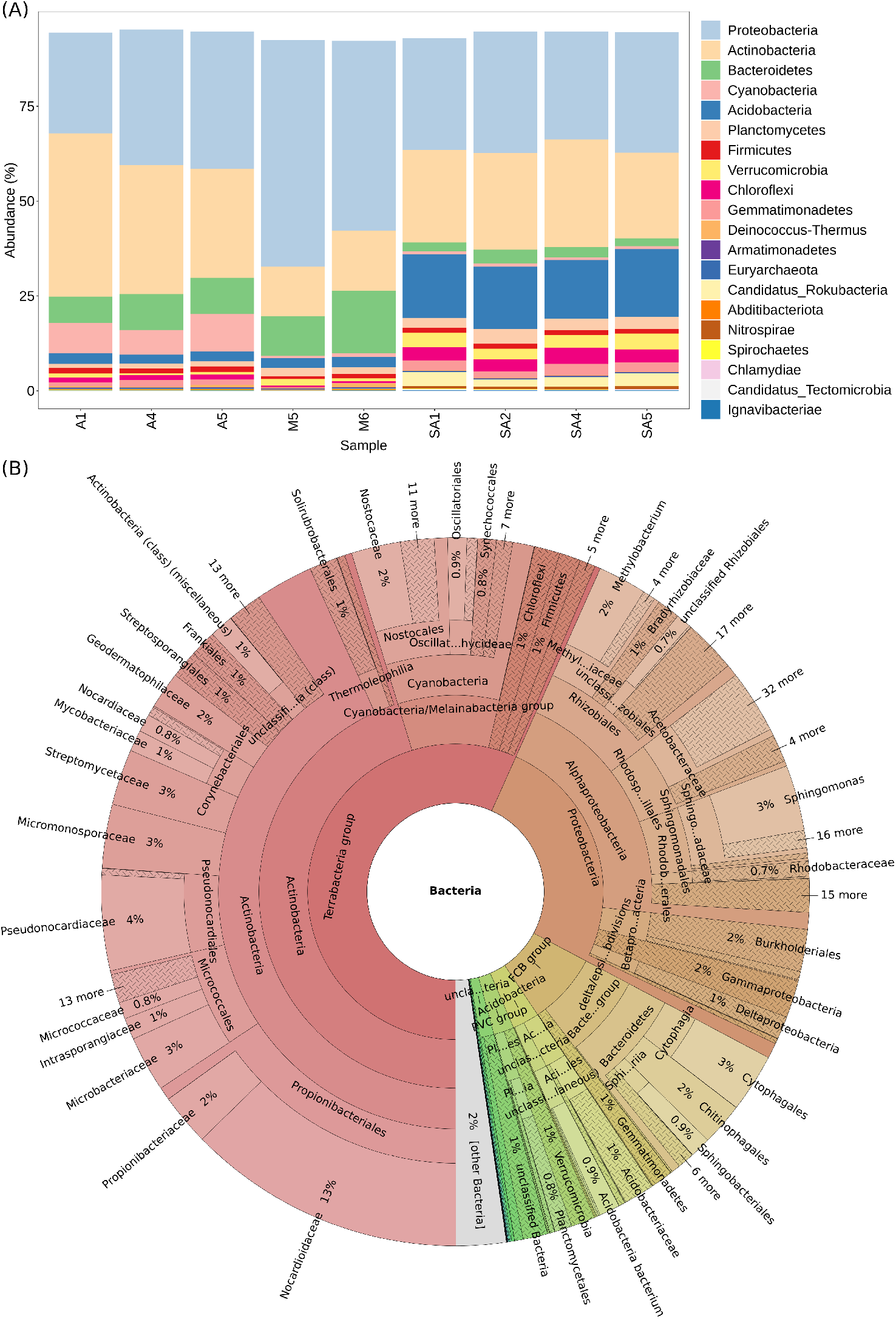
Bacterial diversity in microbial communities of avocado-associated bark and soil. A) Relative abundance in all samples of the 20 phyla that showed the highest abundance in bark. B) General overview of the bacterial diversity in an avocado bark sample from Malinalco (Sample A1). An interactive version of this figure can be found at https://kbase.us/n/69195/32/.

**Figure 3.**
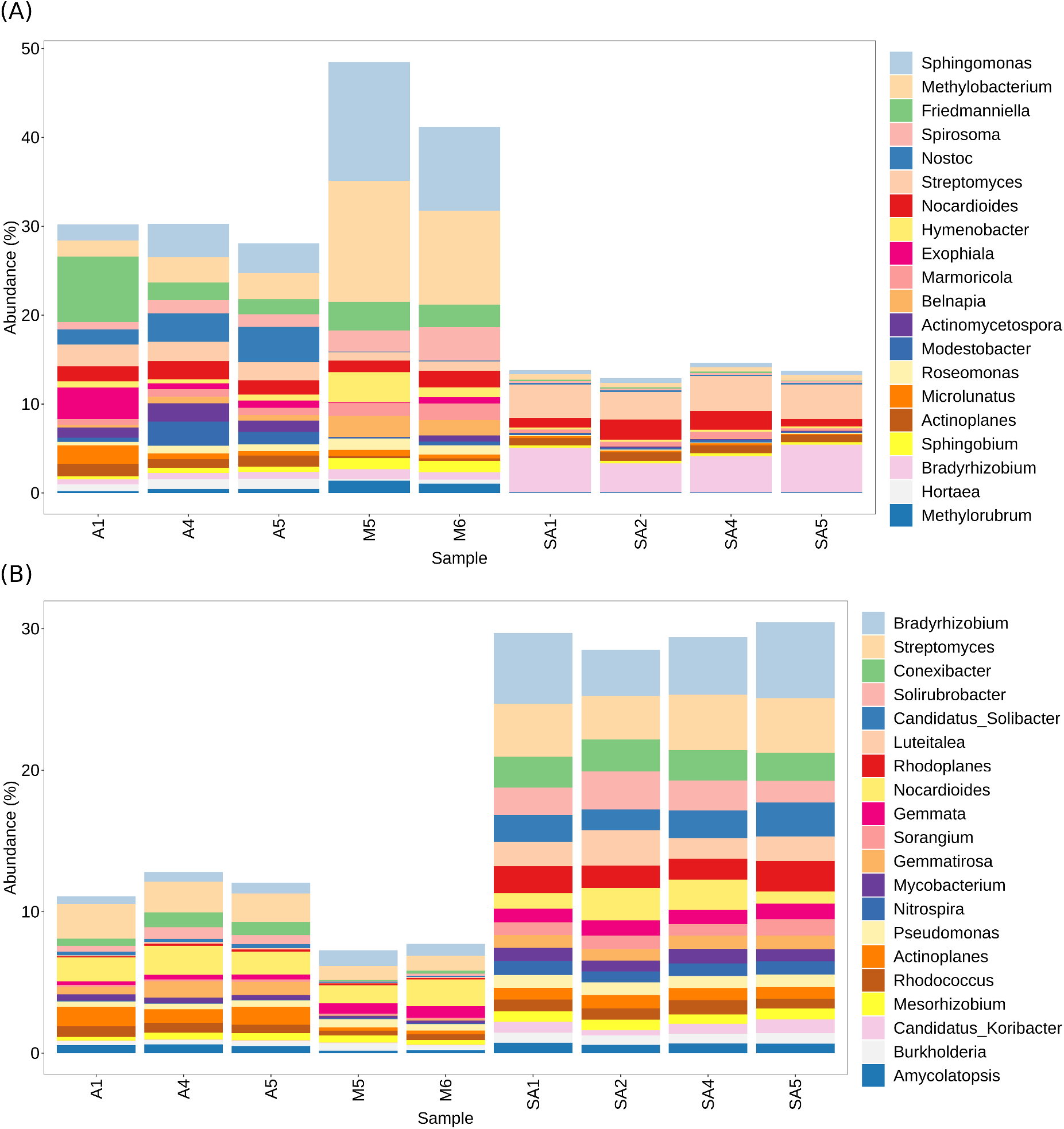
Prokaryotic genus abundances in microbial communities of avocado-associated bark and soil. A) Relative abundance in all samples of the 20 genera that showed the highest abundance in bark. B) Relative abundance in all samples of the 20 genera that showed the highest abundance in rhizospheric soil.

From the phylum Actinobacteria, several genera were among the most abundant in the avocado tree bark environment, including *Friedmanniella, Streptomyces, Nocardioides, Modestobacter, Microlunatus, Pseudonocardia, Conexibacter* and *Rhodococcus.* Bacteroidetes, another dominant phylum, had three genera among the most abundant, namely *Spirosoma, Hymenobacter* and *Flavisolibacter*. Also, the Gemmatimonadetes genus *Gemmatirosa* was one of the most abundant in bark. Interestingly, the cyanobacterium *Nostoc* was highly abundant in the Malinalco bark samples, but not in the Morelia samples.

In rhizospheric soil, the most abundant genus was *Bradyrhizobium* (*Alphaproteobacteria*), followed by the Actinobacteria *Streptomyces* and *Conexibacter,* and the acidobacterium *Luteitalea* (Fig. 3B). All these genera are common inhabitants of soils. Most of the fungi detected in our study belonged to the phylum Ascomycota (Fig. 4, S11-S18). Another phylum, Basidiomycota, was also present in avocado bark, as well as small numbers of other phyla. We note that, as fungi are multicellular, the number of read counts is not a good measure of individual abundance, but rather a general indication of the presence of the group. In rhizospheric soil, the abundance of fungal reads detected was negligible compared to bacteria, perhaps due to the large populations of bacteria found in these environments (Trevors, 2010), which could have overwhelmed fungal reads during sequencing. The large majority of fungi observed in soil samples also belonged to the Ascomycota and Basidiomycota.

**Figure 4.**
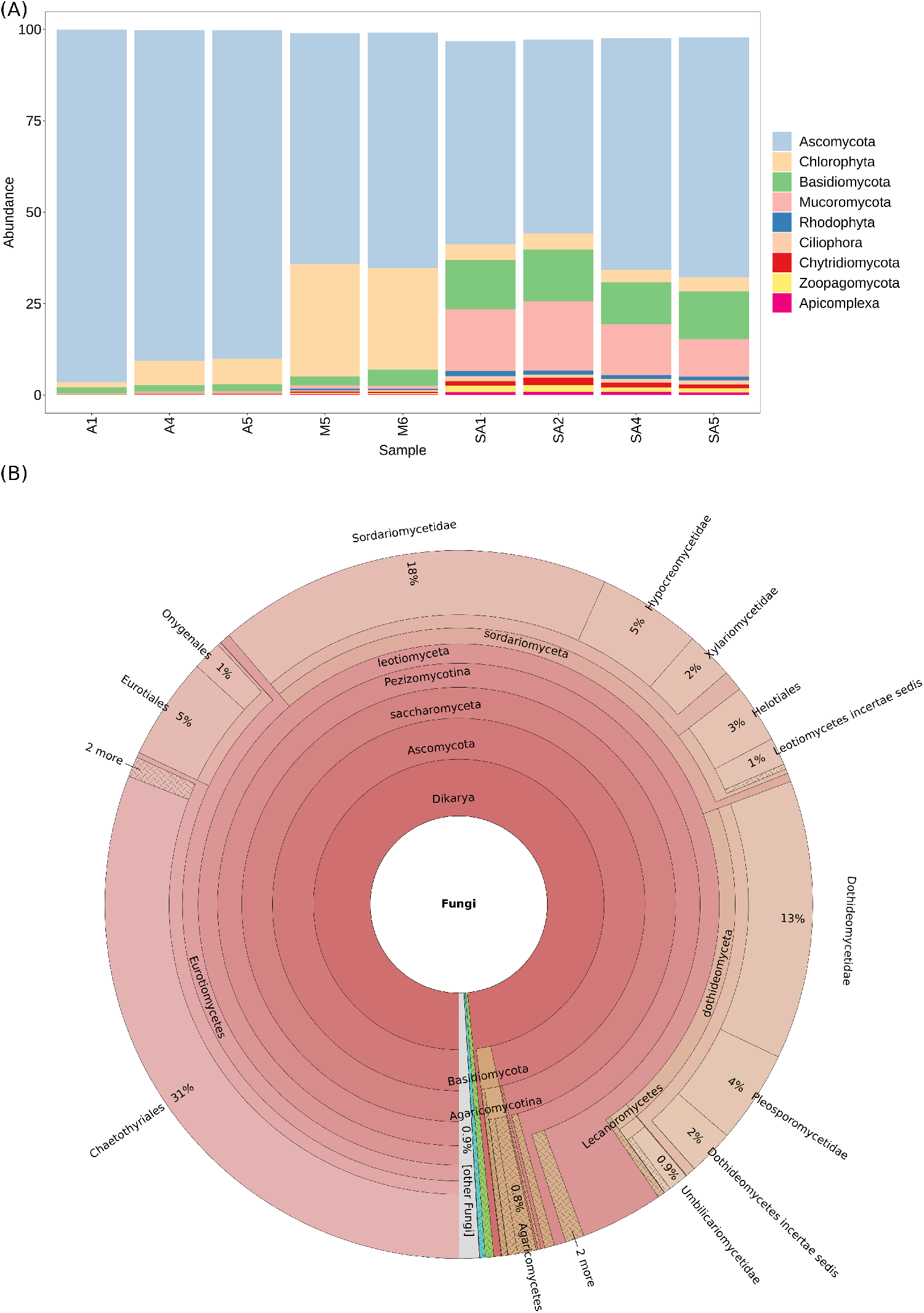
Eukaryotic diversity in microbial communities of avocado-associated bark and soil. A) Relative abundance of the 20 phyla that showed the highest abundance in bark. Abundances were normalized to the total abundance of eukaryotes. B) General overview of the fungal diversity in an avocado bark sample from Malinalco (Sample A1). An interactive version of this figure can be found at https://kbase.us/n/69195/32/.

Abundant fungal genera in avocado bark included *Exophiala, Hortaea, Friedmanniomyces, Aspergillus, Pseudocercospora, Cladophialophora, Baudoinia, Acidomyces, Rachicladosporium, Cercospora, Zymoseptoria, Dothistroma* and *Fusarium* (Fig. 5A)*. Valsa, Umbilicaria* and *Endocarpon* were abundant in the Malinalco samples, but not in Morelia. Conversely, *Paraphaeosphaeria* was very abundant in the Morelia samples, but rare in Malinalco. *Rhizophagus* was the most abundant fungal genus in rhizosferic soil, followed by *Fusarium* and *Aspergillus*. Other fungal genera, like *Penicillium, Cordyceps* and *Hassallia* were very abundant in some of the rhizospheric soil samples, but absent in others.

**Figure 5.**
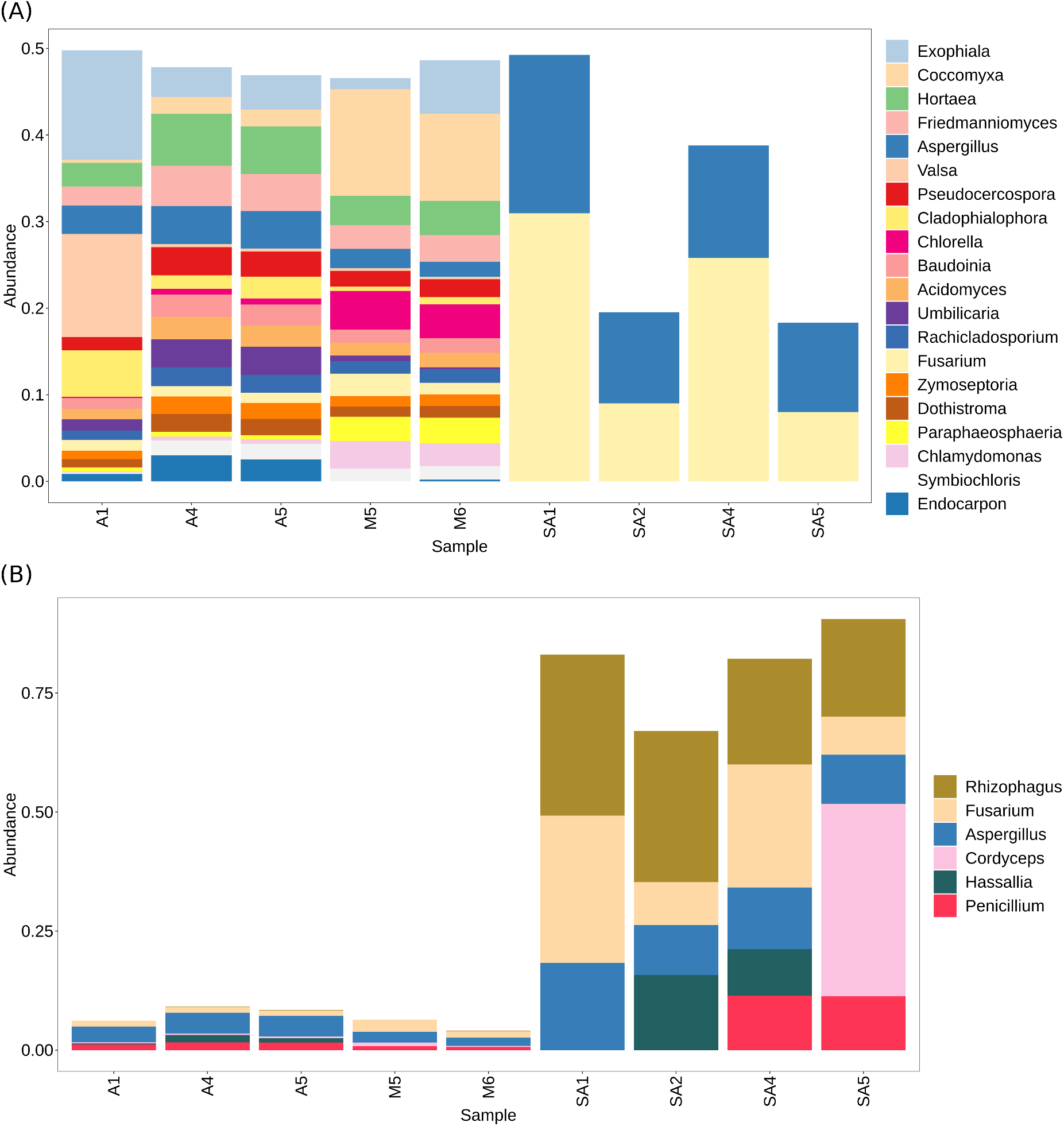
Eukaryotic genus abundances in microbial communities of avocado-associated bark and soil. A) Relative abundance in all samples of the 20 genera that showed the highest abundance in bark. B) Relative abundance in all samples of the 20 genera that showed the highest abundance in rhizospheric soil.

Another group of eukaryots that showed consistent differences between bark and rhizospheric soil samples, independent of bark origin, were the green algae (Fig. S19). For instance, the family *Trebouxiaceae* was observed in all bark samples, but was absent from the rhizospheric soil communities.

Archaea in bark and rhizospheric soil communities were relatively rare, as compared to bacteria. Nevertheless, the consistent presence of certain archaeal groups was observed in avocado barks from both orchards (Fig. 6, S20-S27). The Archaea in bark samples were largely composed of Euryarchaeota, with Thaumarchaeota and Crenarchaeota also present, as well as a small representation of other phyla. In contrast, in soil there were many more Thaumarchaeota present, while the rest of the archaeal phyla had similar abundances.

**Figure 6.**
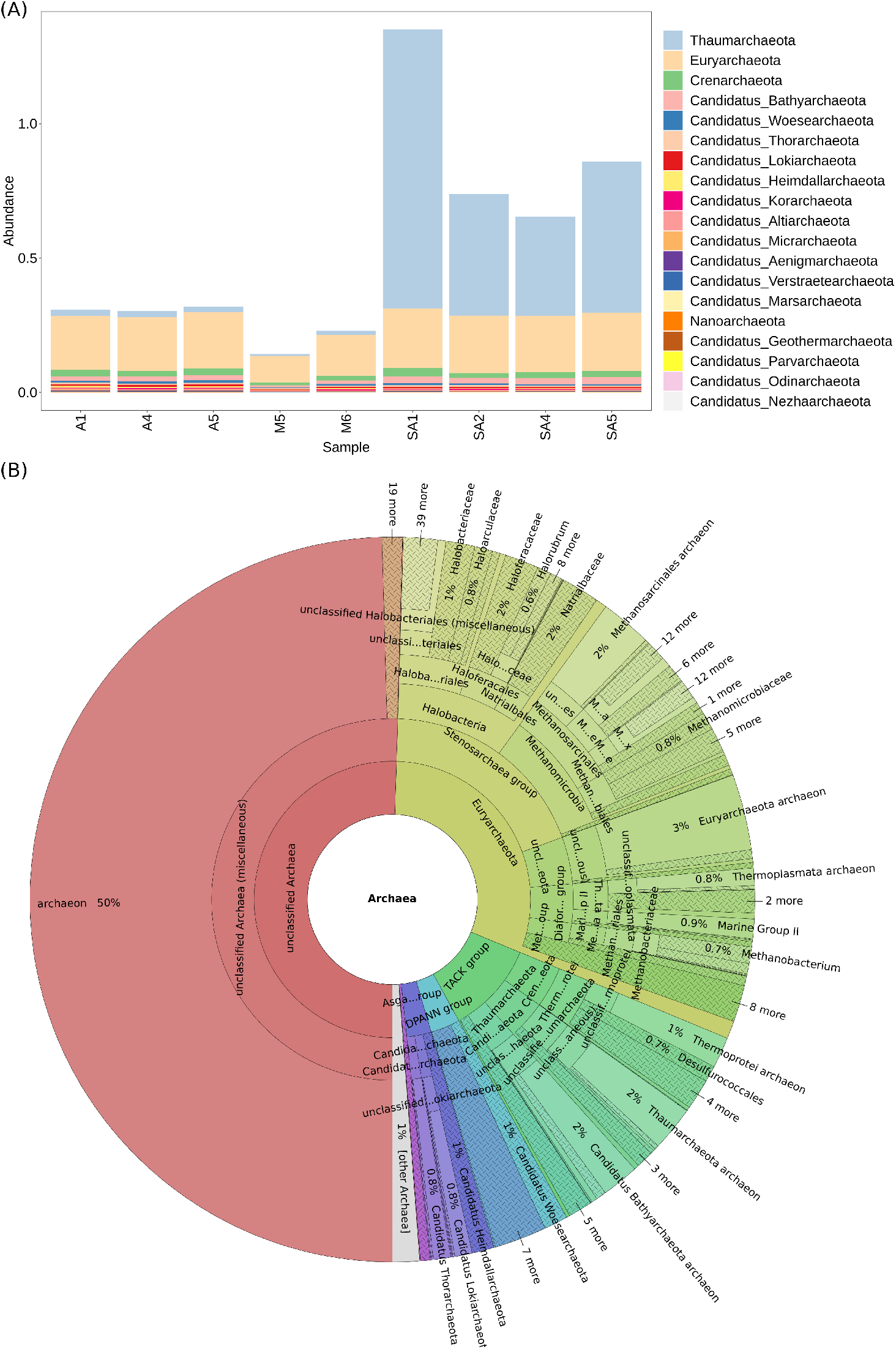
Archaeal diversity in microbial communities of avocado-associated bark and soil. A) Relative abundance of the 20 phyla that showed the highest abundance in bark. B) General overview of the archaeal diversity in an avocado bark sample from Malinalco (Sample A1). An interactive version of this figure can be found at https://kbase.us/n/69195/32/.

The taxonomic composition of non-Taumarchaeota Archaea was strikingly similar between bark and rhizospheric soil samples. For example, the families *Methanomicrobia* and *Methanomada* had very similar proportions of genera for all samples (Fig. S28-S29). This similarity was not observed in all Archaea groups, for example, the TACK group had different patterns of abundance for the bark and rhizospheric soil samples (Fig. S30).

Methanotrophic Archaea were very diverse and abundant in rhizospheric soil, as well as in bark from both orchards, and had strikingly similar taxonomic compositions. Most known orders containing methanogenic Archaea (Vanwonterghem *et al.* 2016) were present, including Methanomicrobiales, Methanosarcinales, Methanomassiliicoccales, Methanobacteriales, Methanococcales, Verstraetearchaeota and Bathyarchaeota. Many Archaea of the order Halobacteriales, which is usually considered an extremophyl group, where also present in all samples.

## Discussion

By sequencing the DNA of avocado bark trees at a whole-community level, we were able to conduct an in-depth metagenomic analysis of the microbial diversity of this environment. Host contamination was negligible, most likely because outer bark consists mostly of dead cells (Franceschi *et al.* 2005), and plant cell walls might be somewhat resistant to the lysis procedures applied. These results were attained by directly inserting small bark pieces into the extraction kits’ first vial, and this procedure is recommended for future studies on bark microbiomes.

By comparing the bark and rhizospheric soil samples from Malinalco, we characterized the microbial communities associated with avocado trees in these two plant compartments. Communities of bark from different trees were clearly more similar between them, compared to the rhizospheric soil samples, as shown by beta-diversity analysis and correlations between abundances of prokaryotic genera (Fig. 1 and S2). This was supported by the relative abundances of specific taxa, including bacteria, archaea, fungi and green algae, which were consistently more similar within either bark or soil (Fig. 2B, 4B, 5B and S3-S30). Furthermore, the Malinalco avocado bark communities were very similar to the Morelia bark communities, providing evidence that the observed community structure is characteristic of avocado bark as a microbial habitat. The importance of the plant compartment as a major determinant of microbial community composition has been previously reported for other plant species, such as *Agave* spp. (Coleman-Derr *et al.* 2015), vine (Vitulo *et al.* 2019) or tomato (Allard *et al.* 2016). Our results extend these findings to the bark of living trees, and show that the bark of avocado is a well-defined microbial environment, favoring the establishment of particular microbial groups. As many of the microorganisms found in bark were also present in soil (Fig. 1D), the latter could act as a reservoir for the aerial parts of plants (Martins *et al.* 2013; Grady *et al.* 2019).

Even though the microbial communities from the bark samples from Malinalco and Morelia had highly correlated genus abundances, and were similar in the abundance of many taxonomic groups, they also had important differences, and could be well distinguished based on Bray-Curtis distances. This illustrates that variation can be present in bark environments, when different geographic regions, management practices, times of the year, and tree age, among other factors, are in play, as shown for other plant species (Vitulo *et al.* 2019; Arrigoni *et al.* 2020). The similarities between the microbial communities of bark from the Malinalco and Morelia orchards, suggest there could exist a “core bark microbiome” from avocado trees. However, research in more areas, and with larger sample sizes would be needed to confirm this hypothesis.

There is limited information on microbial communities of bark from living trees. In this work, avocado tree bark bacterial communities were dominated by representatives of the phyla Actinobacteria, Proteobacteria (in particular, Alphaproteobacteria) and Bacteroidetes, consistently with previous findings in other woody plants, including apple, pear, *Ginkgo biloba,* wild trees from the Atlantic Rainforest in Brazil, and grapevine (Lambais *et al.* 2014; Leff *et al.* 2015; Arrigoni *et al.* 2018; Vitulo *et al.* 2019; Arrigoni *et al.* 2020). Other phyla observed among the dominant bark microbial communities included Cyanobacteria (only in the Malinalco orchard in this study), Deinococcus-Thremus (in apple and pear), Acidobacteria (in *Ginkgo bioloba*) and Flavobacteria (in wild trees from the Atlantic Rainforest). Thus, bark microbial communities are consistently dominated by the same major phyla, with some variation observed in particular cases. This consistency shows that the bark environment is a defined habitat for specific groups of bacteria, as opposed to a surface that can be colonizated by random bacterial groups. This is similar to the relative stability at the phylum level observed in soil communities, where a few phyla tend to dominate as well (Janssen, 2006; Aguirre-von-Wobeser *et al.* 2018; Delgado-Baquerizo *et al.* 2018).

Interestingly, some of the most abundant bacterial genera found in bark in this study, including *Hymenobacter* and *Sphingomonas,* were also among the most abundant in 16S rRNA gene surveys in other trees (Leff *et al.* 2015; Arrigoni *et al.* 2018; Vitulo *et al.* 2019; Arrigoni *et al.* 2020). Furthermore, strains of *Streptomyces* and *Arthrobacter*, two bacterial genera which were very abundant in this study, were isolated from bark of Hass avocado (Dunlap *et al.* 2017), suggesting a consistent presence of some microbes in bark communities even at the genus level.

*Sphingomonas,* the most abundant genus found in bark at both orchards, is very common on plant surfaces, including bark (Leff *et al.* 2015; Arrigoni *et al.* 2018; Vitulo *et al.* 2019; Arrigoni *et al.* 2020) and leaves (Rivas *et al.* 2004; Bodenhausen *et al.* 2014; Cha *et al.* 2019; Delmotte *et al.* 2019). *Sphingomonas* can grow on many recalcitrant substrates (White *et al.* 1996), which might be advantageous for living heterotrophically on bark organic materials. Furthermore, it can produce complex exopolysaccharides (White *et al.* 1996), which could aid in water retention on plant surfaces. This genus has both phytopahtogenic (Buonaurio *et al.* 2001; Deldavleh *et al.* 2013; Kini *et al.* 2017) and plant-beneficial strains, which provide protection against pathogens (Berg and Ballin, 1994; Innerebner *et al.* 2011) and abiotic stress (Halo *et al.* 2015; Asaf *et al.* 2017). These complex relationships of *Sphingomonas* with plants are an interesting case for future studies on the evolution of plant-microorganism interactions.

*Methylobacterium*, the second most abundant genus found in avocado bark microbial communities, is a facultative methylotroph (Vorholt, 2012), which might benefit from both plant (Wang *et al.* 2016) and microbial methane emission, as many potentially methane-producing Archaea were present in the community. Interestingly, *Methylobacterium* isolates were also obtained from the phyllosphere of two *Lauraceae* species (the same plant family as *P. americana*) in Mexico, and displayed antifungal activity *in vitro* against *Fusarium solani* (Báez-Vallejo *et al.* 2020). The dominance of *Methylobacterium* at the bark level could therefore play a potential role in the biocontrol of *Fusarium* dieback in avocado trees.

The cyanobacterium *Nostoc* was the third most abundant genus in bark overall, but found mainly in the Malinalco orchard. The ability of this cyanobacterium to form complex extracellular polysaccharide matrices, together with its ability to fix atmospheric nitrogen (Otero and Vincenzini, 2003), seem to be advantageous adaptations to retain water and obtain nitrogen in the bark environment (Vorholt, 2012; Stone *et al.* 2018). Another very abundant genus in bark was *Spirosoma,* which is commonly found in soils (*e.g.* Joo *et al.* 2017; Li *et al.* 2018), and some species can live as endophytes (Fries *et al.* 2013).

*Friedmanniella* (and closely related *Microlunatus),* another very abundant actinobacterium in bark, is known to accumulate polyphosphate inside their cells (MasZenan *et al.* 1999), which can be used as a phosphorus and energy source (Nakamura *et al.* 1995). Monopolizing nutrients and exploiting alternative energy sources could be an advantageous strategy in this oligotrophic environment.

A large diversity of *Streptomyces* was observed, which are well known for their metabolic diversity and production of useful secondary metabolites, including compounds with antifungal activity (Getha *et al.* 2002; Jog *et al.* 2014; Faheem *et al.* 2015), as well as their capacity to solubilize phosphates (Jog *et al.* 2014).

Some of the most abundant fungi identified in bark are extremophiles, including *Acidomyces, Friedmanniomyces* and *Hortaea,* although they have been found in very different extreme conditions (Selbmann *et al.* 2017; Buzzini *et al.* 2018; Chan *et al.* 2018). It is not clear whether the stress resistance mechanisms of these fungi could be protective in the bark environment. *Umbilicaria* and *Endocarpon* are commonly found forming lichens (Park *et al.* 2016; Rodríguez-Caballero *et al.* 2018), and their presence in the Malinalco samples, where cyanobacteria were also abundant, suggests these symbiotic associations might take place in bark from this orchard, although no macroscopic lichens were visible. Phytopathogenic fungal genera such as *Fusarium, Rachicladosporium, Pseudocercospora, Zymoseptoria, Cercospora, Dothistroma,* and *Aspergillus* were also detected in avocado bark tree from both orchards. The presence of many phytopathogenic fungi in relatively high abundances in bark was surprising, as all the trees appeared healthy and were productive. As not all phytopathogens attack all plant species, and phytopathogenicity can be species or strain specific, the relationship between these microorganisms and the plant cannot be inferred in our study. It is possible that a balance exists in the microbial community, where plant beneficial microorganisms keep phytopathogen populations controlled (Arrigoni *et al.* 2018).

The taxonomic composition of Archaea in avocado bark and soil showed intriguing patterns. First, Thaumarchaeota, the dominant archaeal phylum in rhizospheric soil in this and other studies (*e.g.* Jiao et al. 2019), were almost absent in the bark communities. In contrast, within individual groups of Archaea there were strong similarities between bark and soil. To our knowledge, there are no ecological models to explain such a pattern, and this illustrates our little knowledge on the ecology of Archaea, which should be expanded to have a more complete picture of the microbial world. We can conclude that the microbial communities of avocado bark have a particular, well defined community structure, related to the soil microorganisms, but with their own characteristics in terms of taxonomic composition. Therefore, our results show that the bark environment provides ecological niches for specific taxonomic groups. The study of the metabolic functions present in the avocado bark microbial communities, using our metagenomic dataset, will provide further insight into the adaptations to the bark environment.

This study provides a baseline for the understanding of avocado bark microbial communities, and could guide future efforts in their manipulation for agronomic purposes, as researchers could select microbial groups of interest to follow during manipulation experiments.

## Funding

This work was supported by a grant from the Mexican Council of Science and Technology (CONACyT) and the Mexican Ministry of Education (SEP) [grant number CB-2014-01-242956].

## Conflicts of interest

The authors declare no conflicts of interest

## Acknowledgments

We express our gratitude to the owners of the orchards were the samples were collected, Ulrike Wiegel Torres (Malinalco), Jesús Torres Rodríguez (Malinalco) and José Pedro Rangel Hernández (Morelia), and the caretaker José Guadalupe Arellano Lara (Malinalco). We thank Oliver Aguirre Morales and Emiliano Aguirre Morales for assistance during sampling, and Zachary Crockett, Miriam Land and Elisha M. Wood-Charlson from KBase for support during data analysis and Mayra de la Torre Martínez and Jorge Rocha-Estrada for useful comments on the manuscript.

## Notes

https://kbase.us/n/69195/32/

### Competing Interest Statement

The authors have declared no competing interest.

