## Supplementary materials for "Bark from avocado trees of different geographic locations have consistent microbial communities"

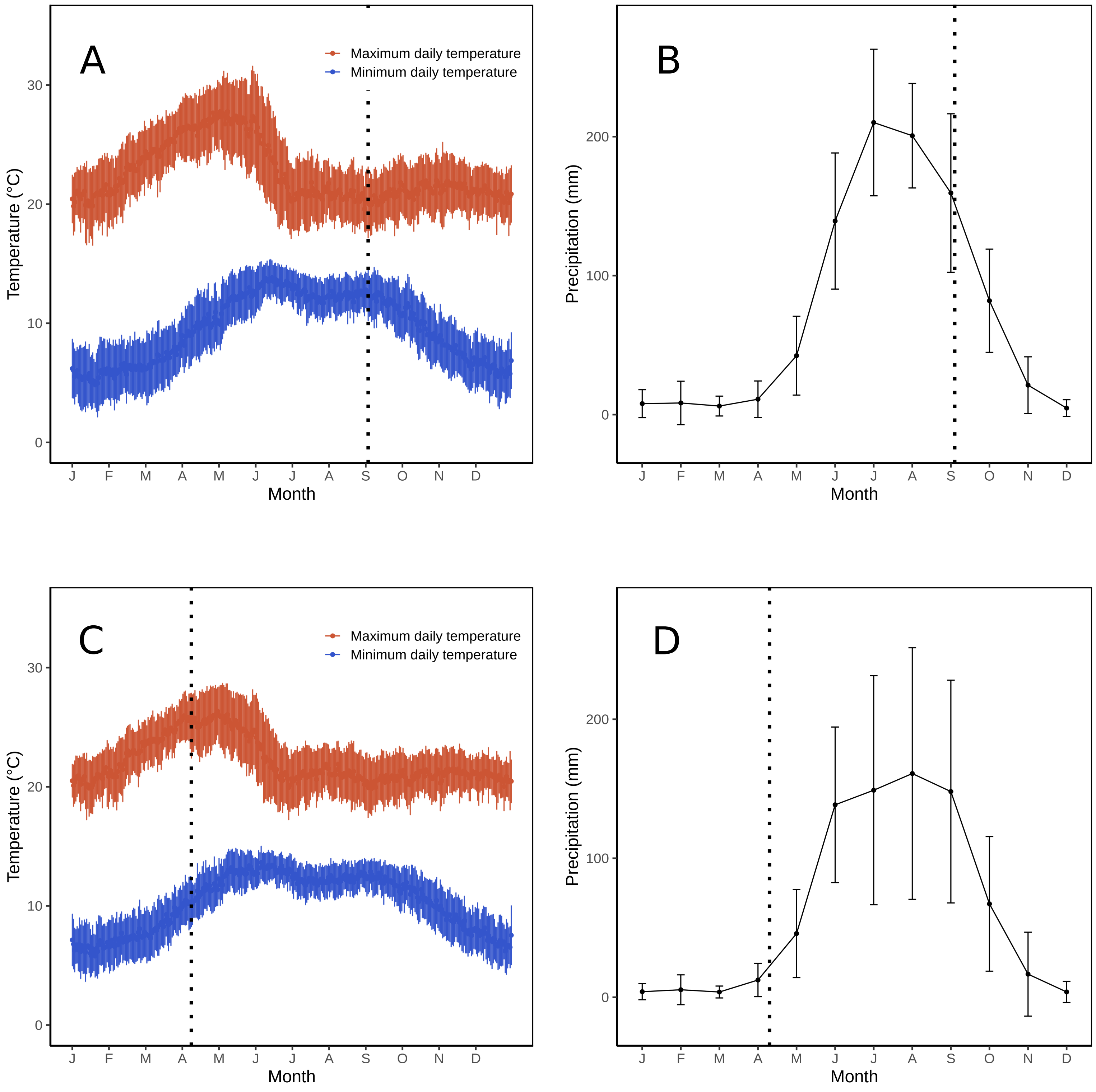


Figure S1. Climate at the orchards of Morelia and Malinalco. A) Maximum and minimum daily temperature at the Morelia orchard, B) Precipitation at the Morelia orchard. C) Maximum and minimum daily temperature at the Malinalco orchard, D) Precipitation at the Malinalco orchard. Data points are means, from 1979 to 2013, and error bars are standard deviations. Precipitation data were calculated monthly from daily data. Vertical dotted lines indicate sampling dates for both orchards.

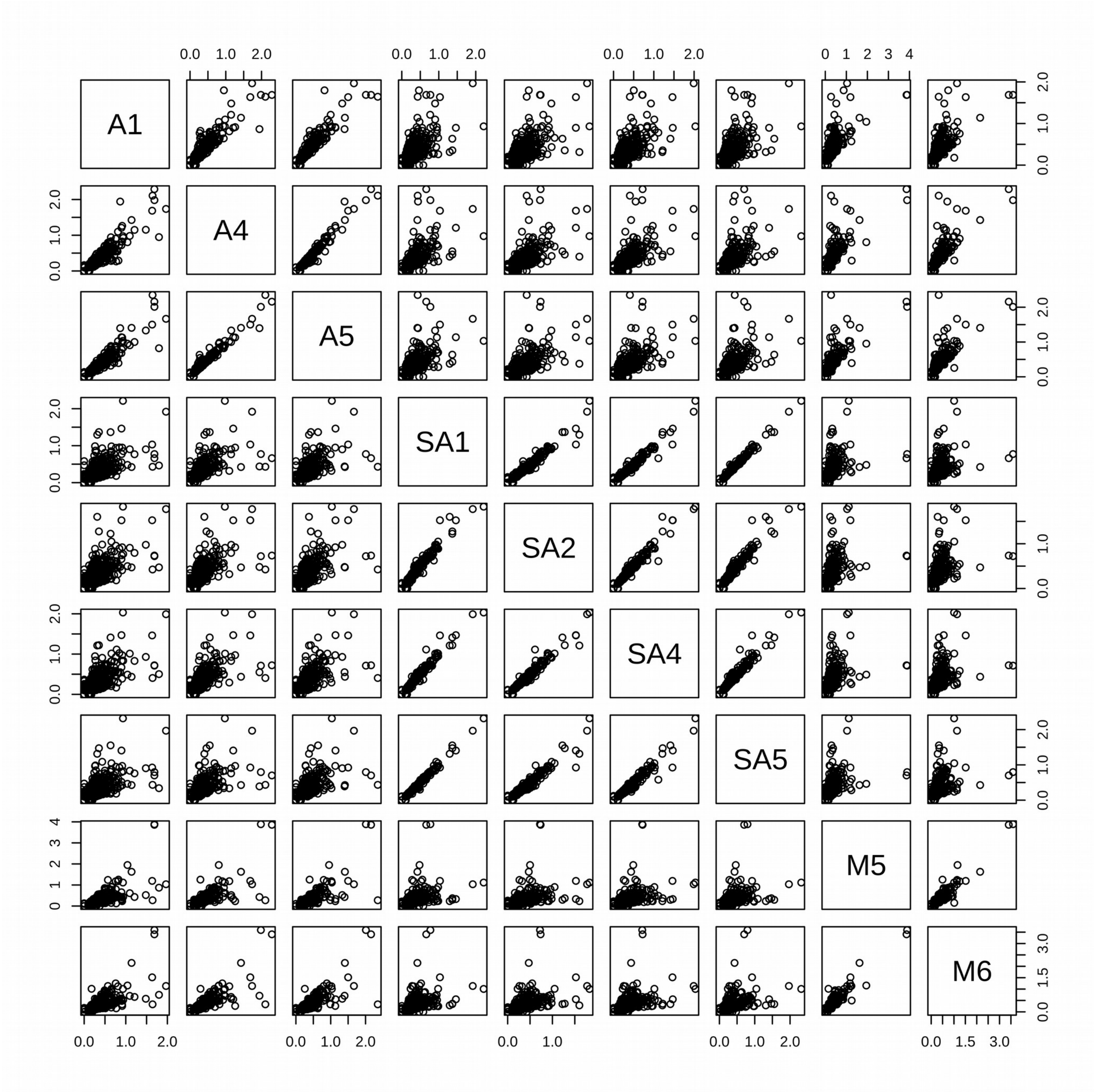


Figure S2. Pair-wise comparisons of the genera abundances of all samples sequenced.

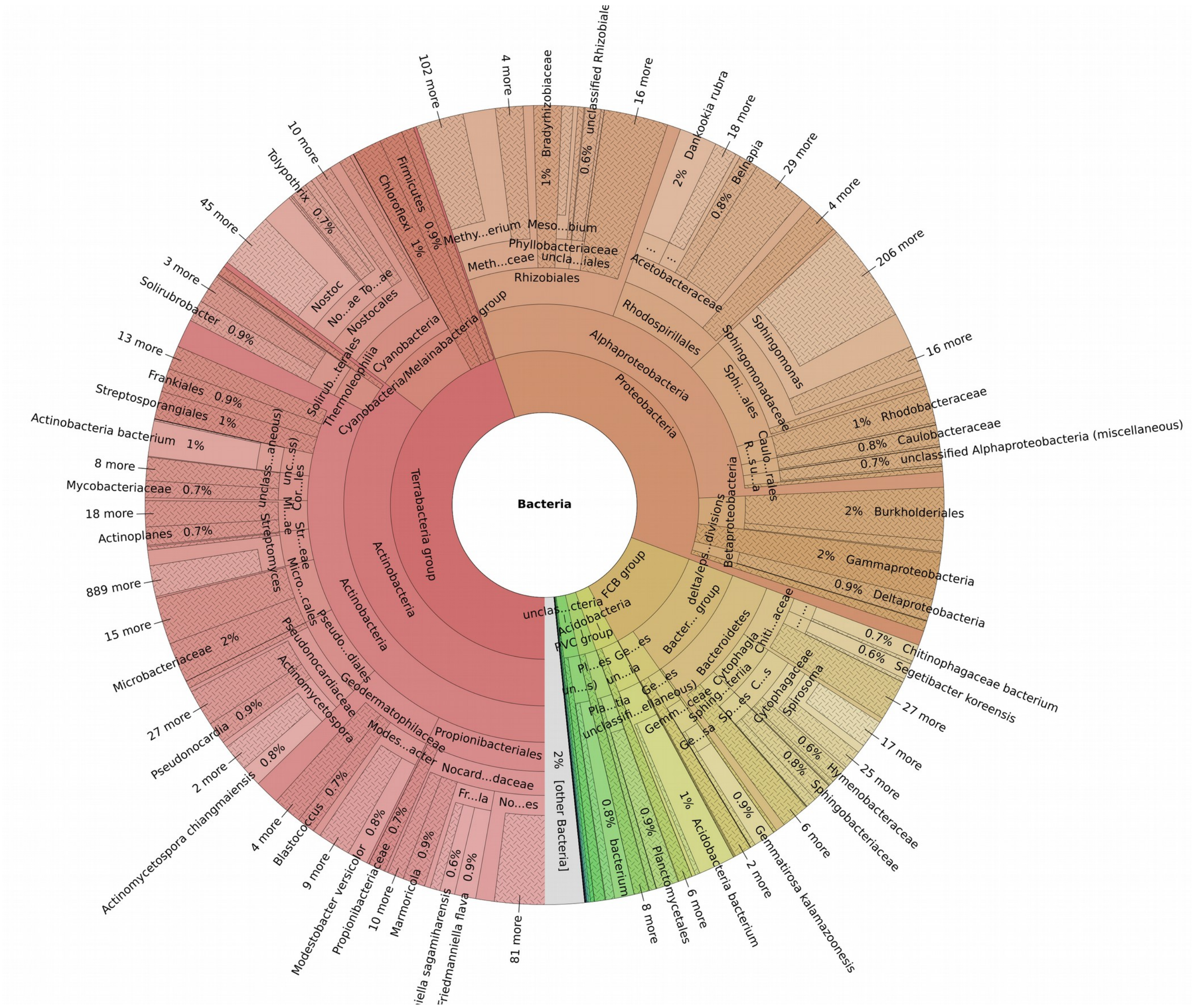


Figure S3. General overview of the bacterial diversity in an avocado bark sample from Malinalco (Sample A4).

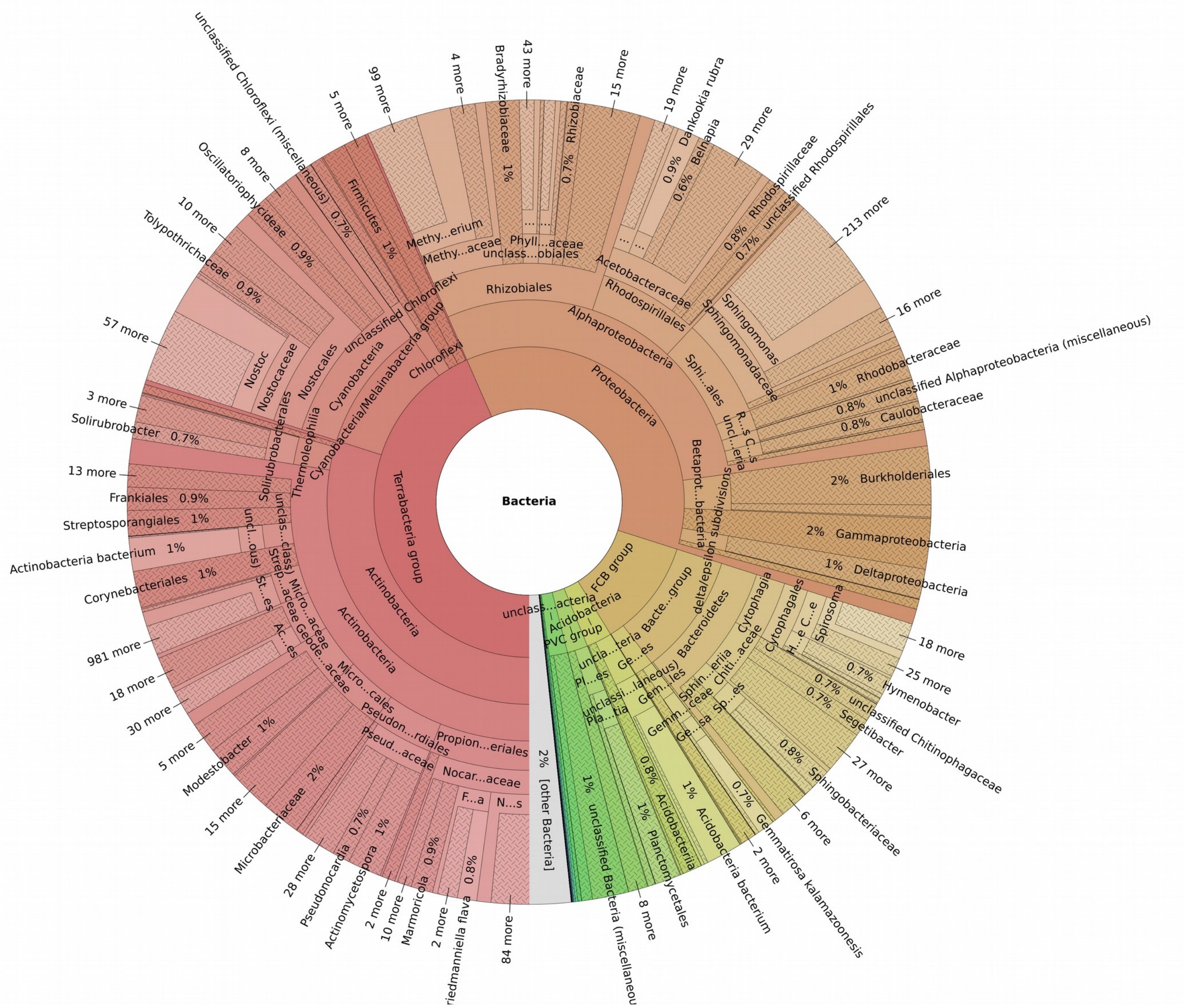


Figure S4. General overview of the bacterial diversity in an avocado bark sample from Malinalco (Sample A5).

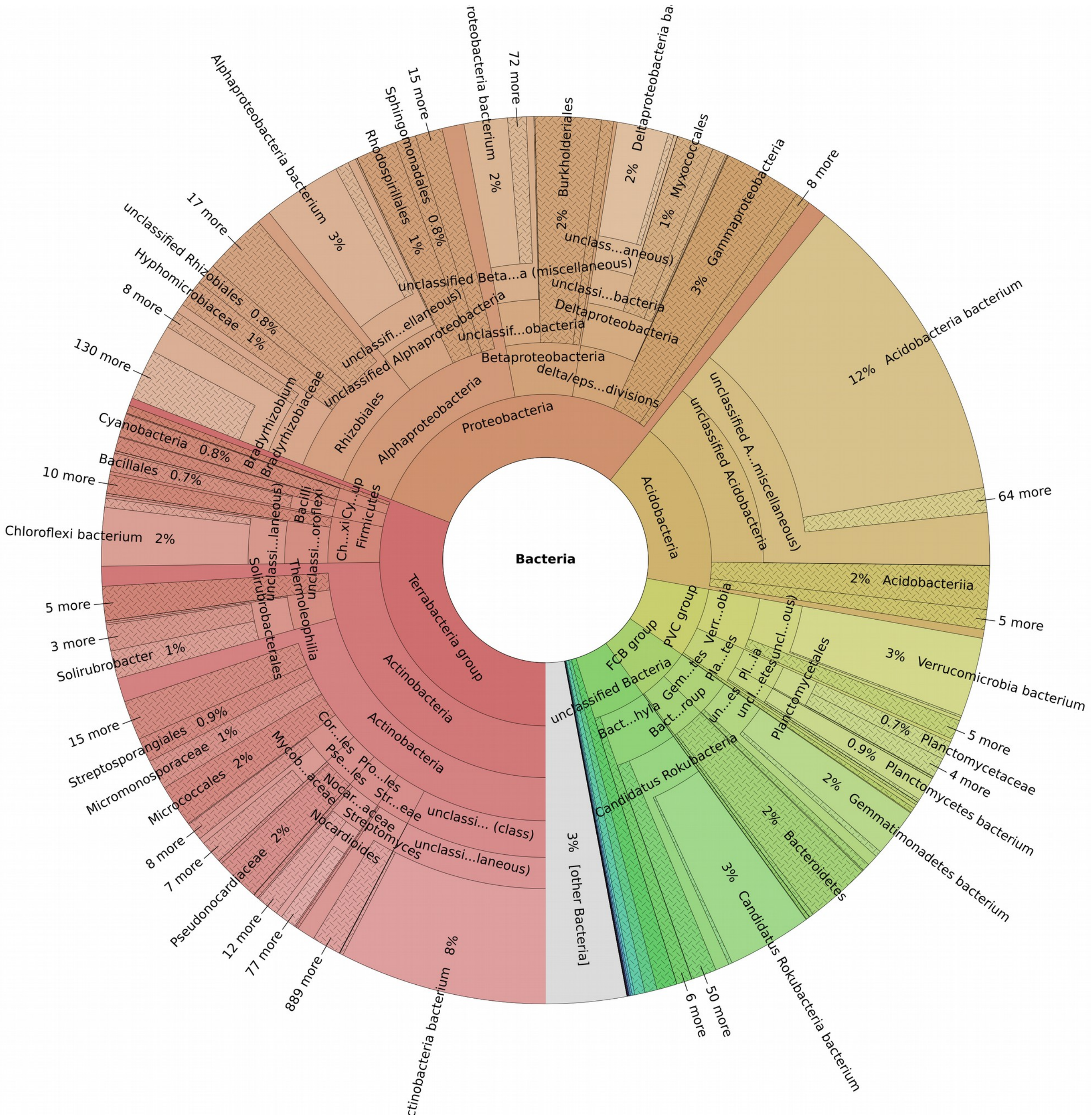


Figure S5. General overview of the bacterial diversity in a soil sample from Malinalco (Sample SA1).

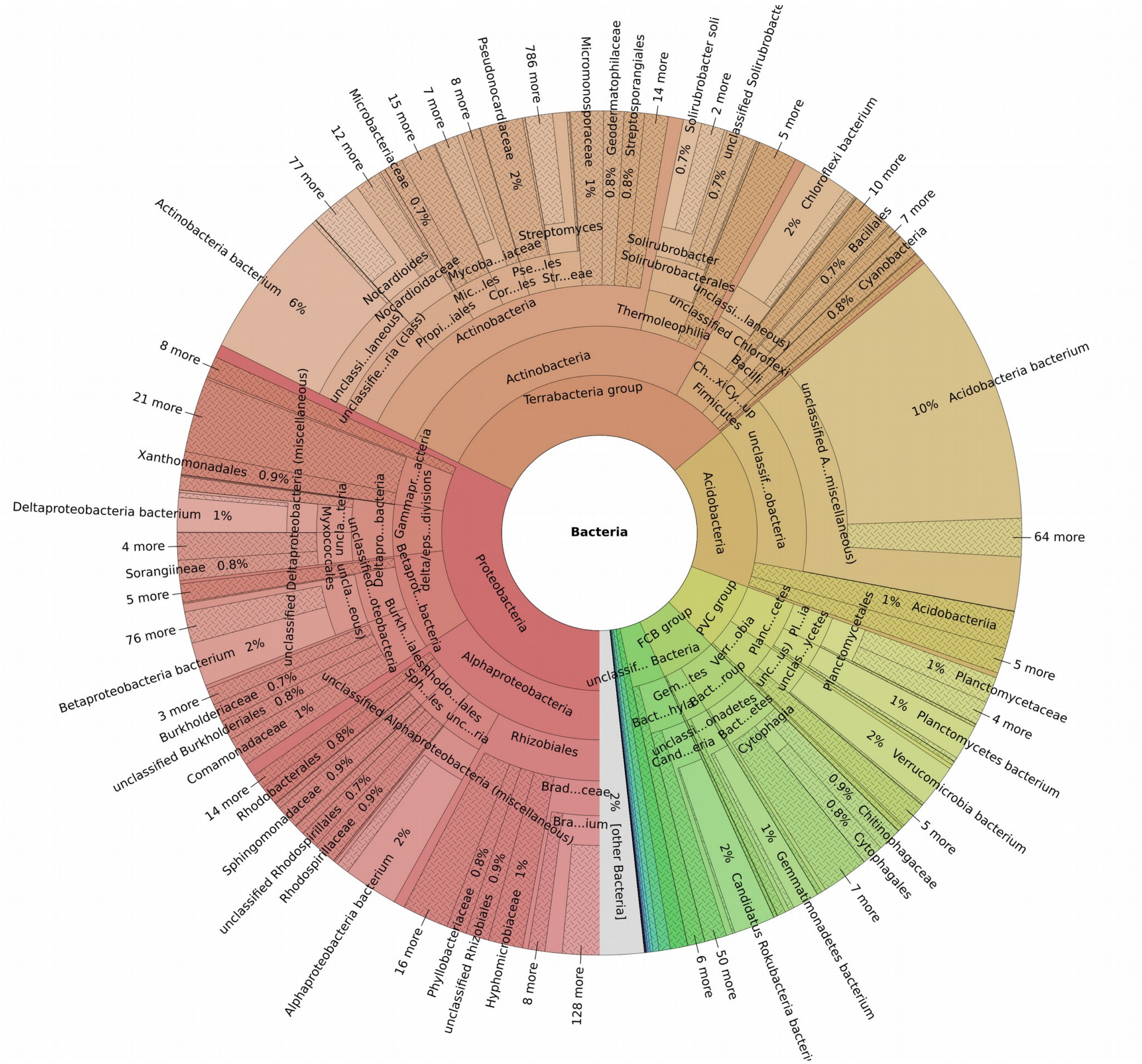

Figure S6. General overview of the bacterial diversity in a soil sample from Malinalco (Sample SA2).

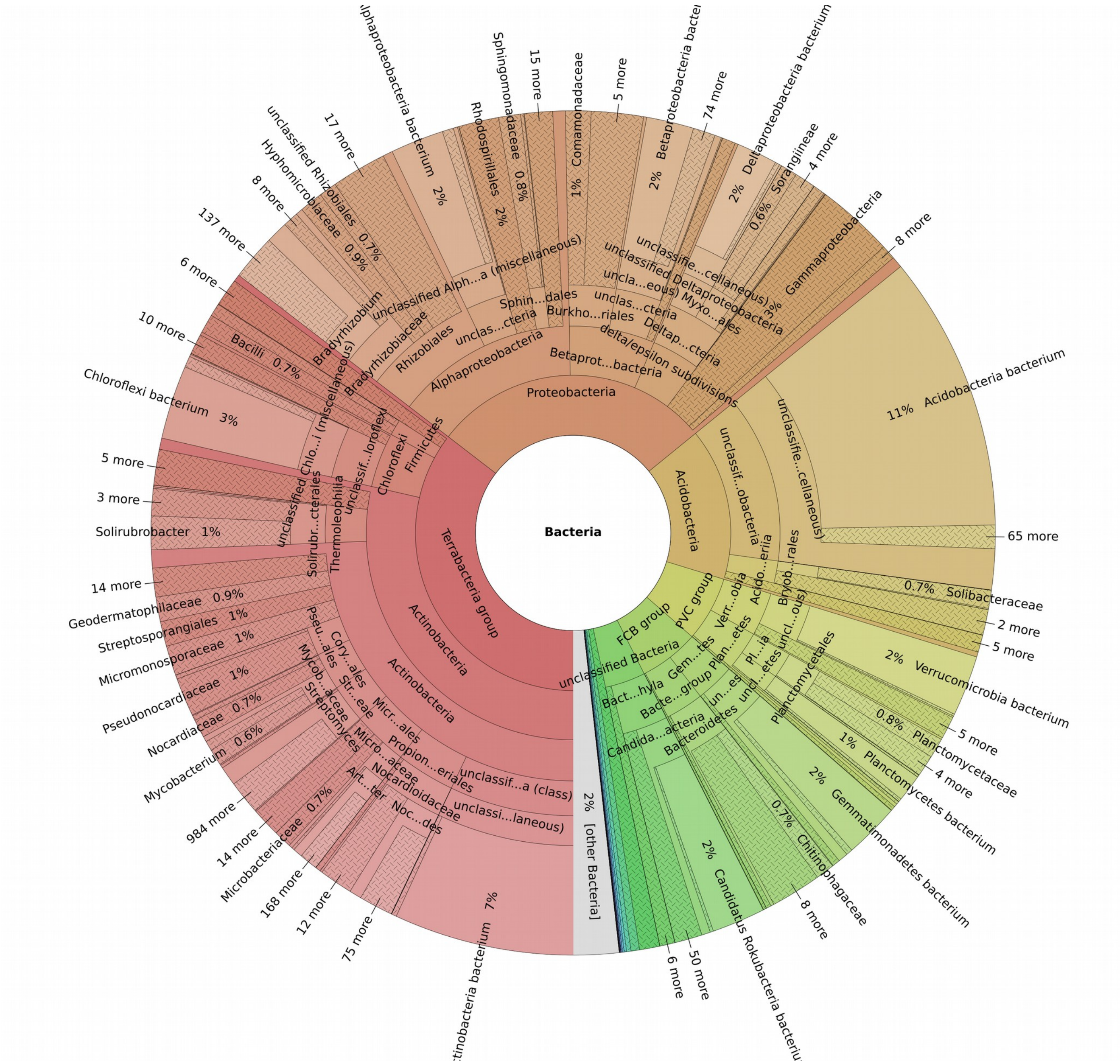

Figure S7. General overview of the bacterial diversity in a soil sample from Malinalco (Sample SA4).

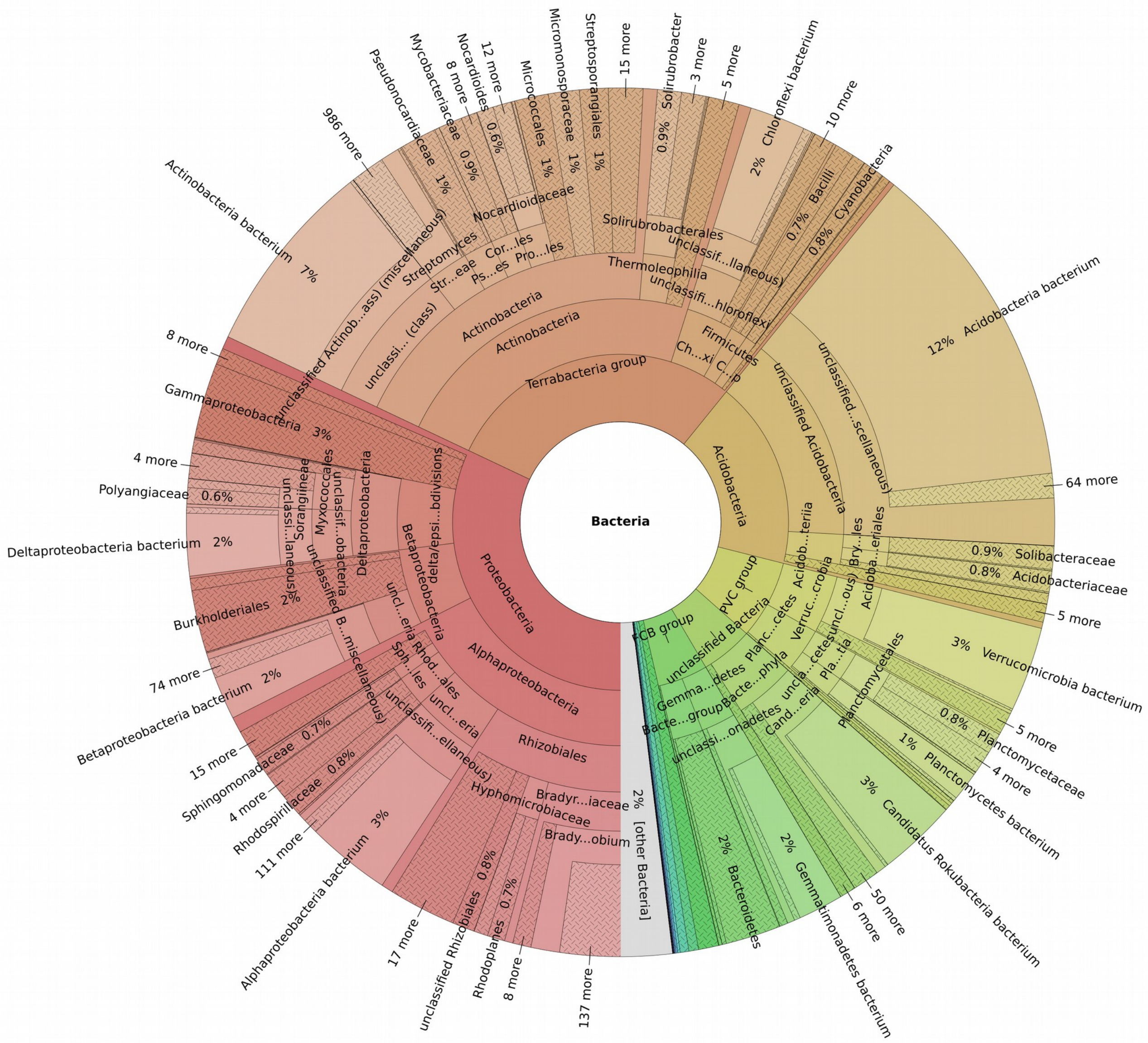

Figure S8. General overview of the bacterial diversity in a soil sample from Malinalco (Sample SA5).

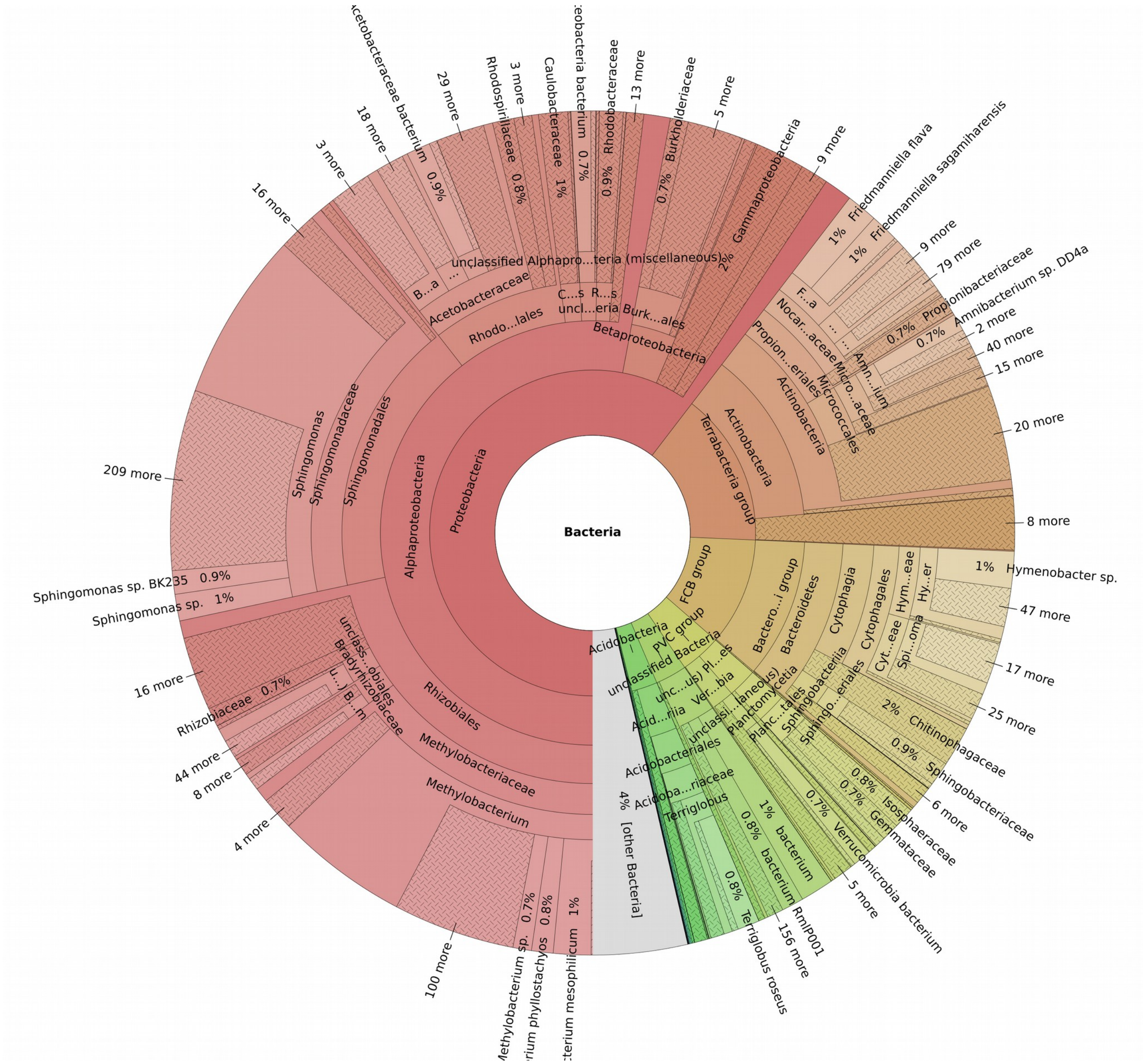


Figure S9. General overview of the bacterial diversity in an avocado bark sample from Morelia (Sample M5).

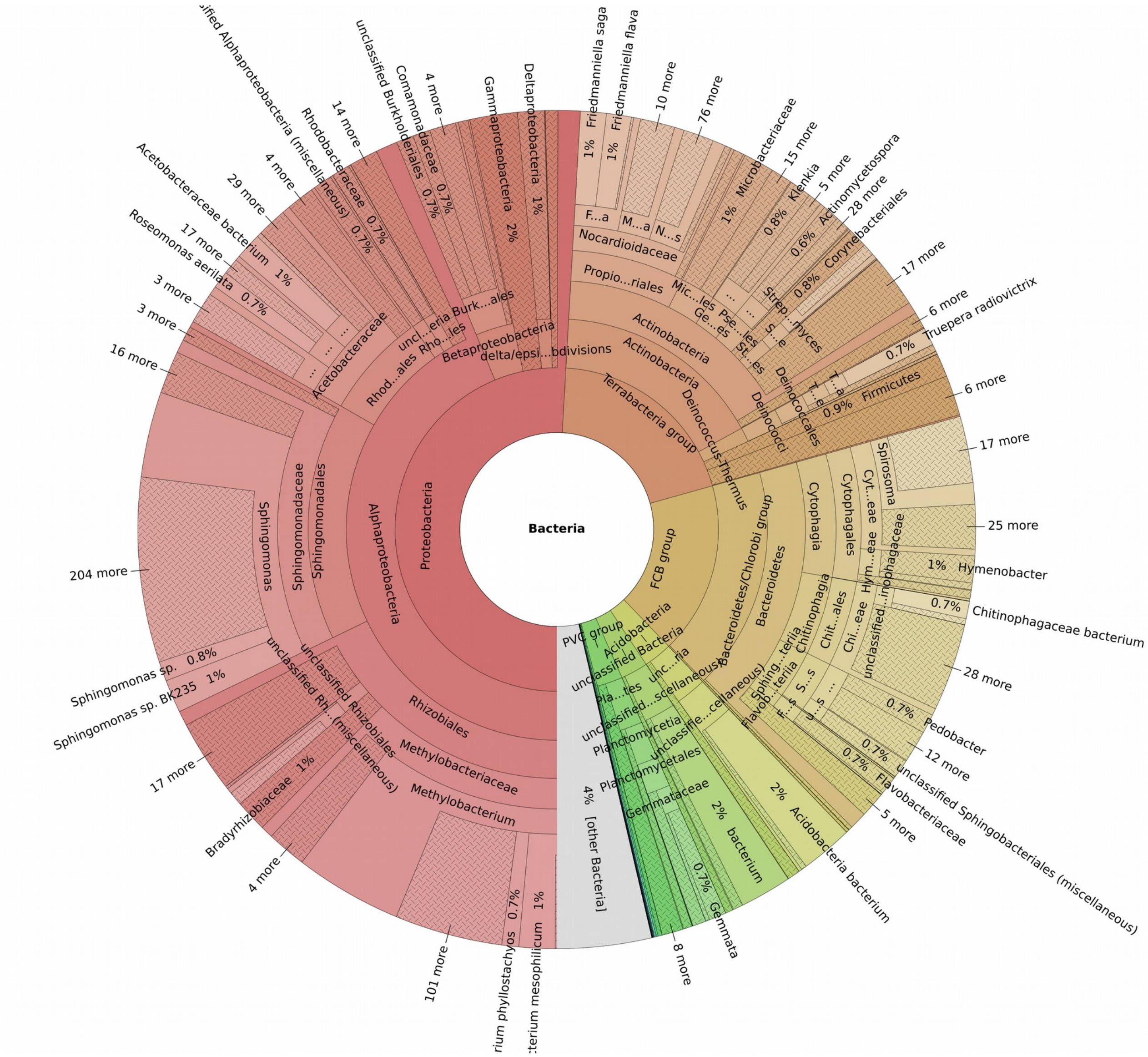

Figure S10. General overview of the bacterial diversity in an avocado bark sample from Morelia (Sample M6).

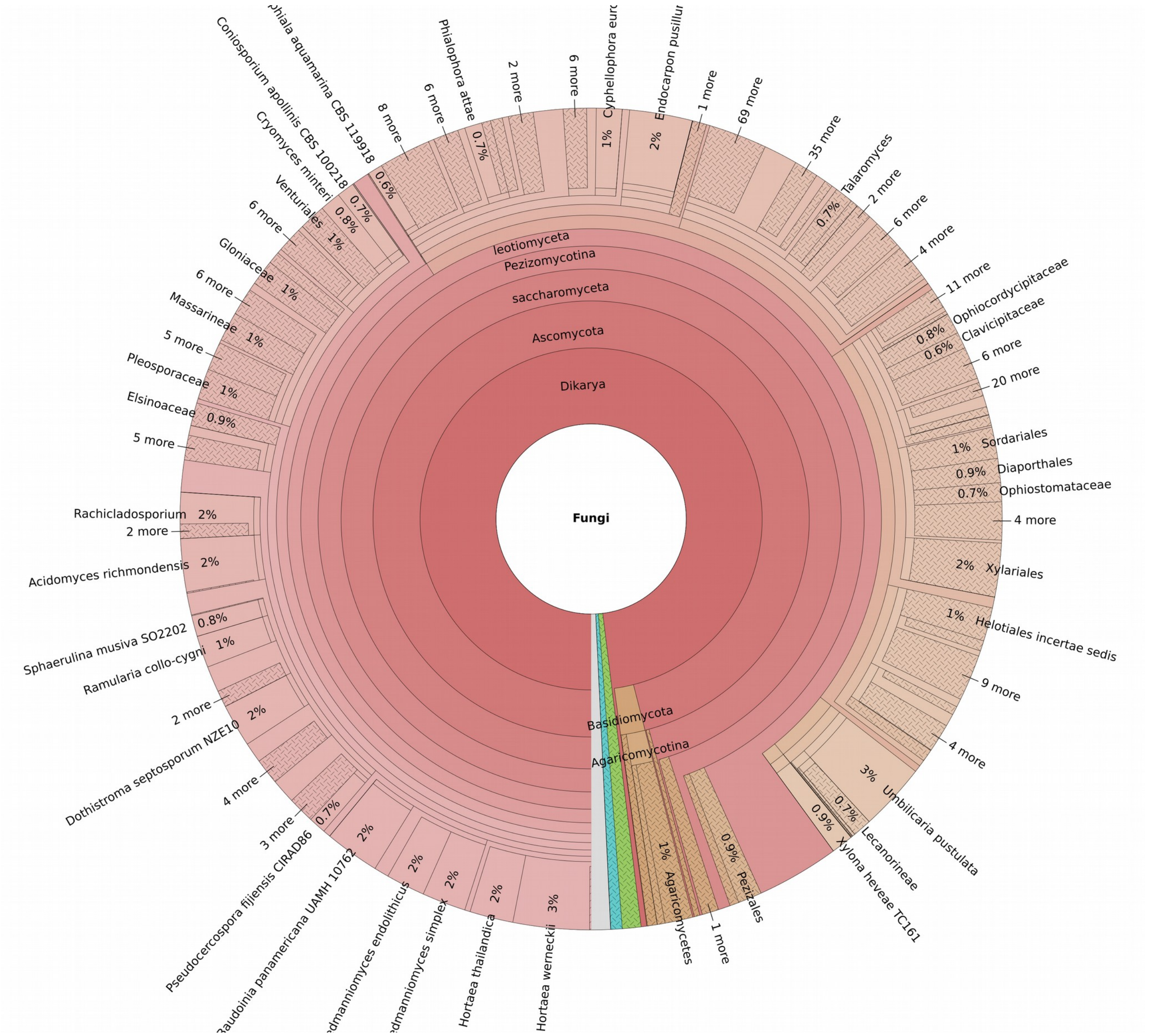


Figure S11. General overview of the fungal diversity in an avocado bark sample from Malinalco (Sample A4).

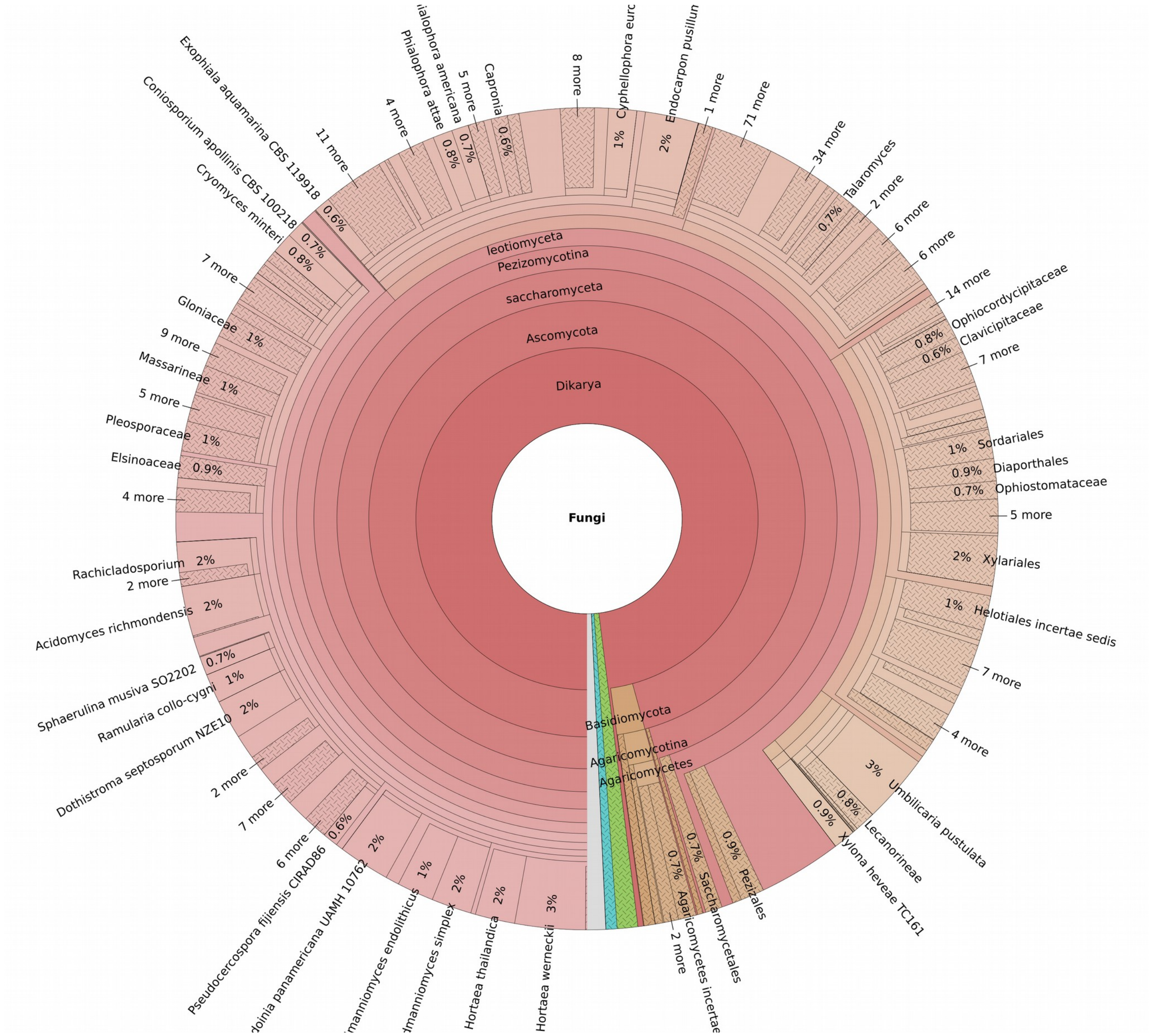


Figure S12. General overview of the fungal diversity in an avocado bark sample from Malinalco (Sample A5).

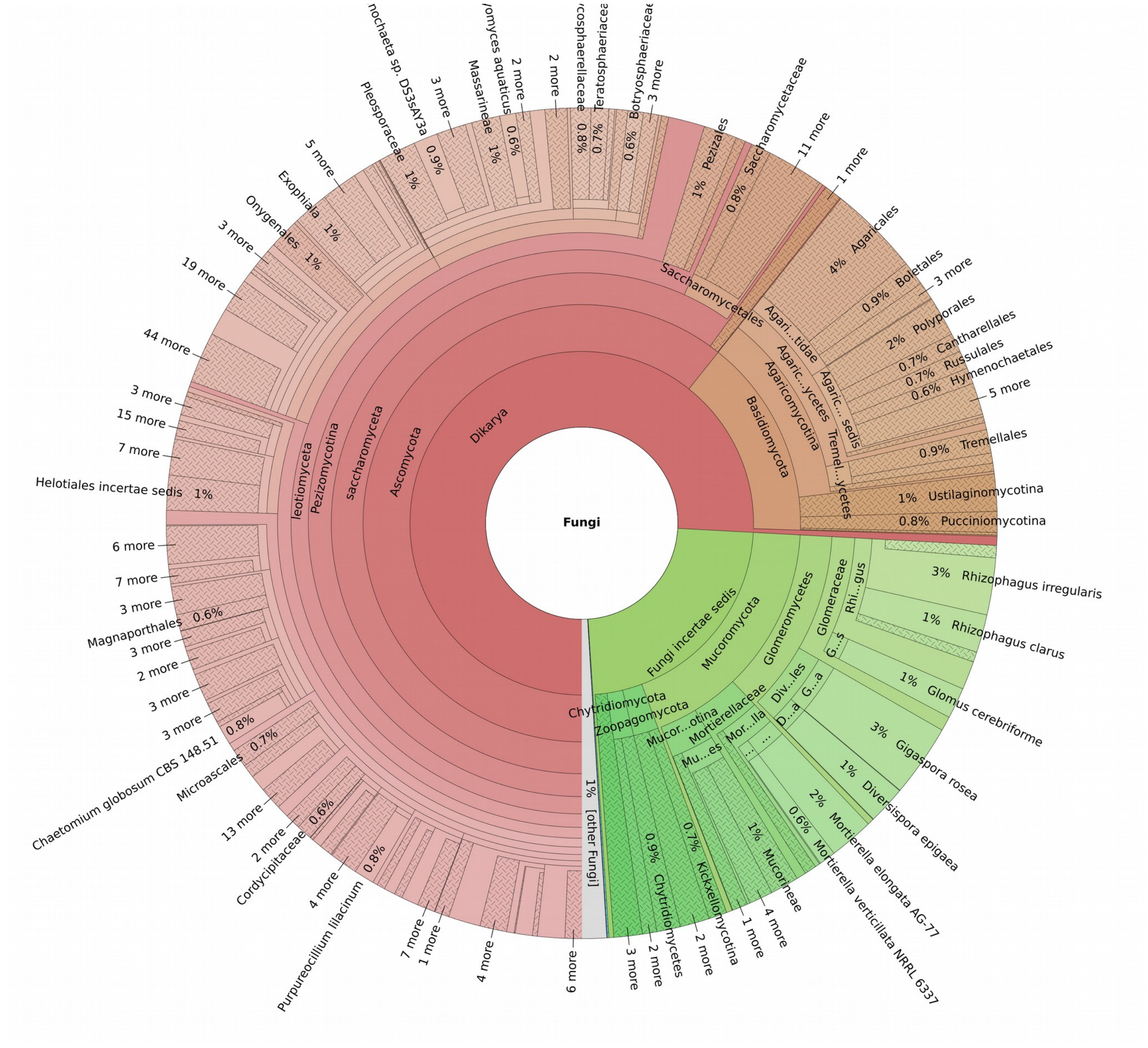

Figure S13. General overview of the fungal diversity in a soil sample from Malinalco (Sample SA1).

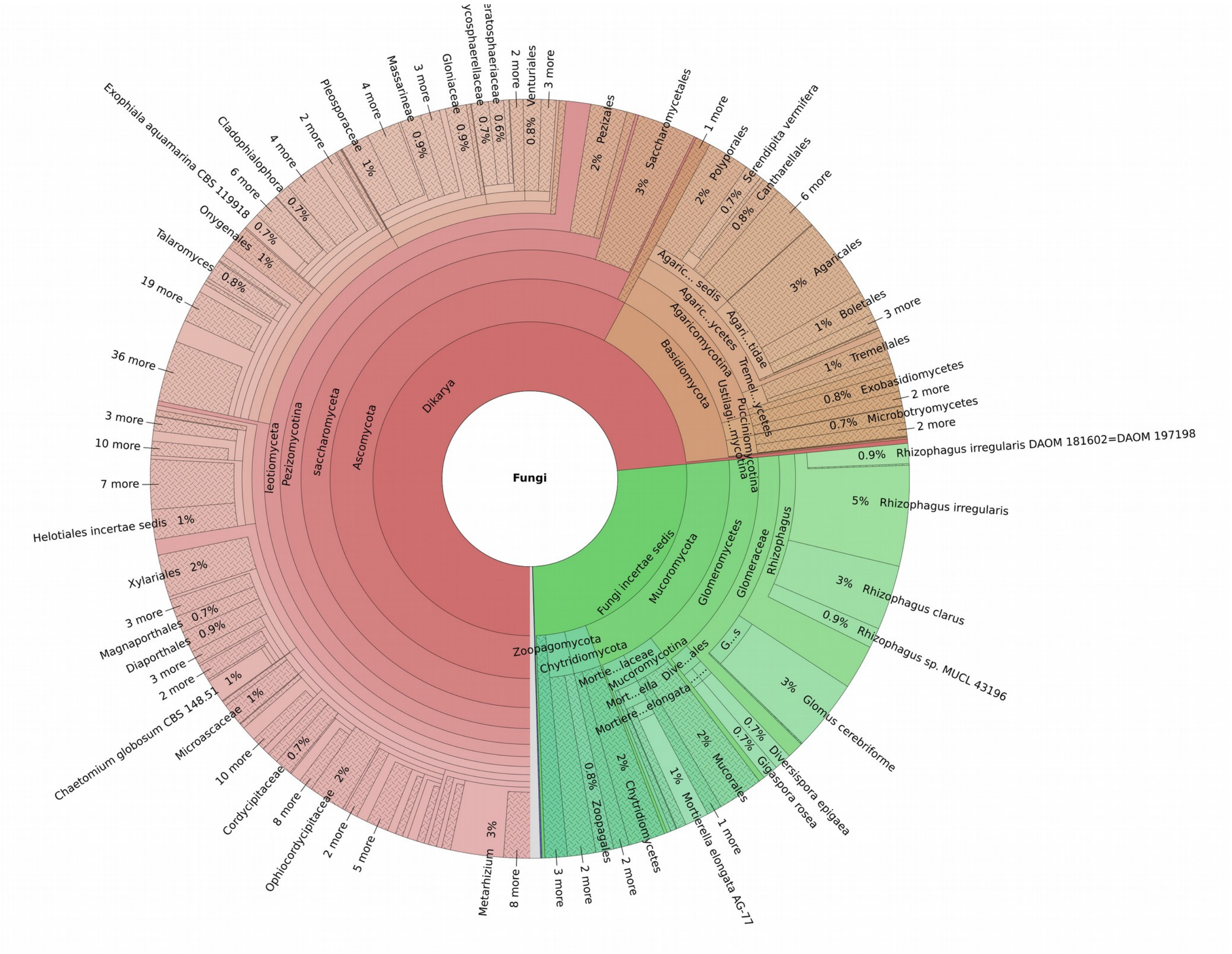

Figure S14. General overview of the fungal diversity in a soil sample from Malinalco (Sample SA2).

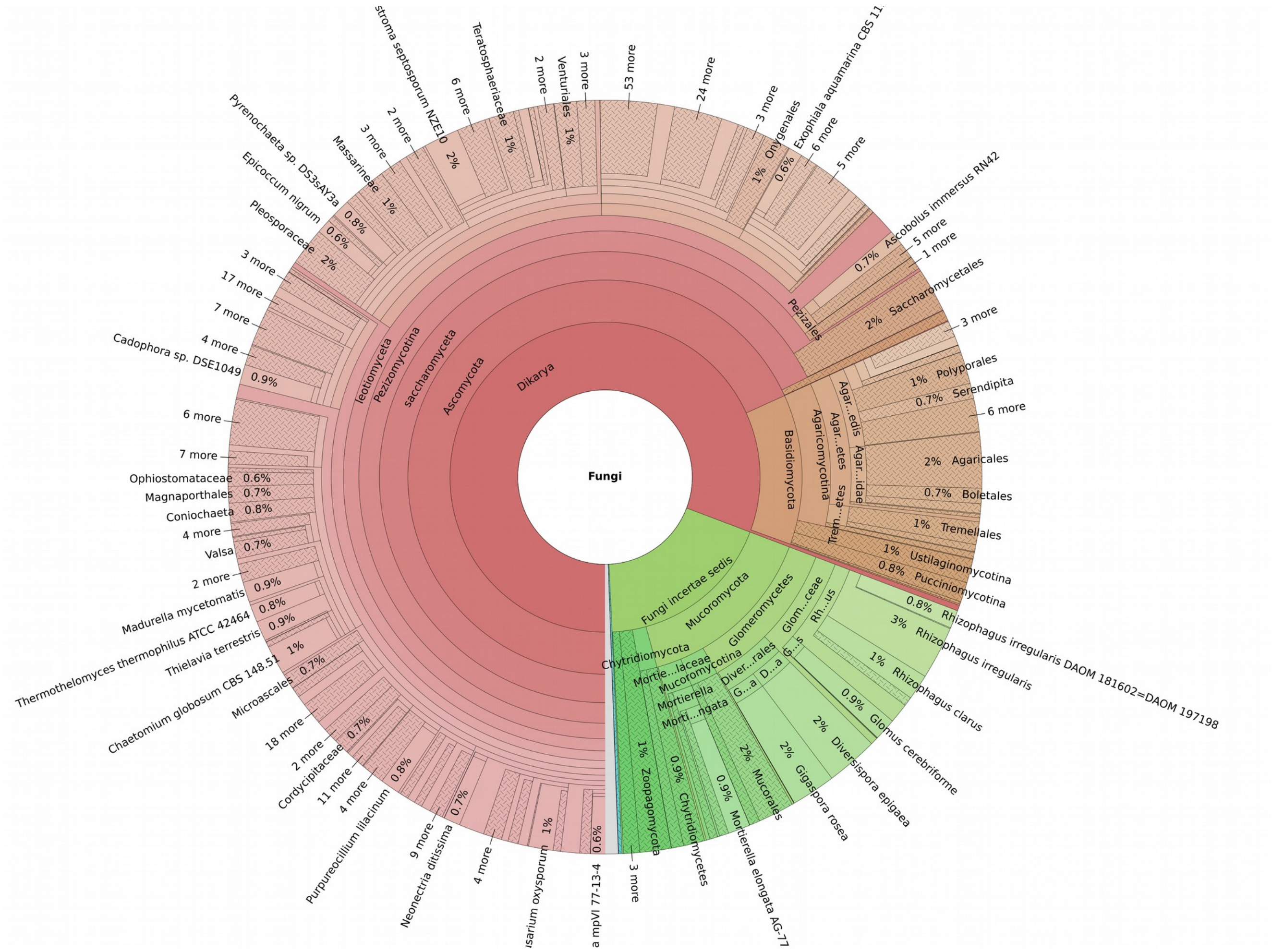

Figure S15. General overview of the fungal diversity in a soil sample from Malinalco (Sample SA4).

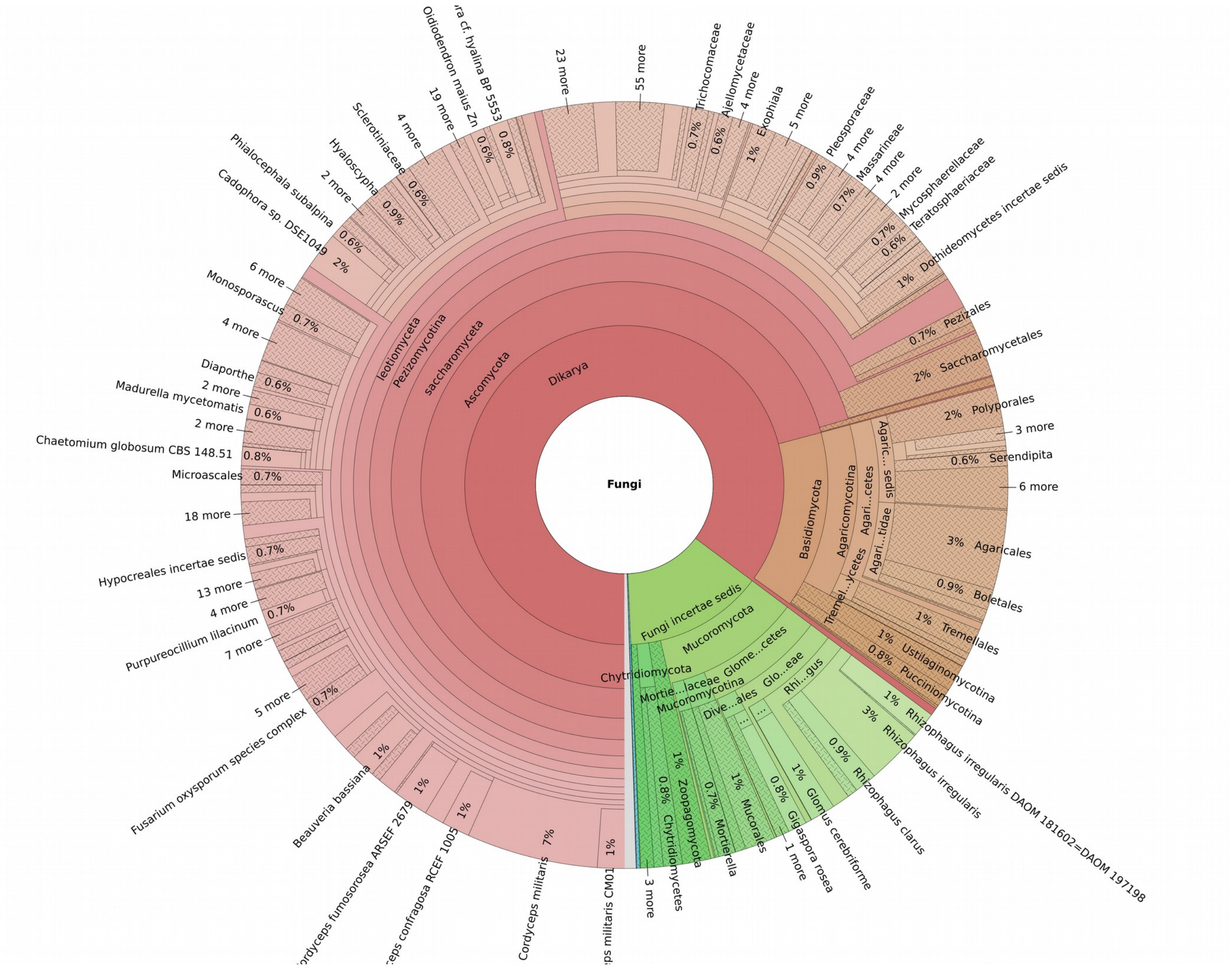


Figure S16. General overview of the fungal diversity in a soil sample from Malinalco (Sample SA5).

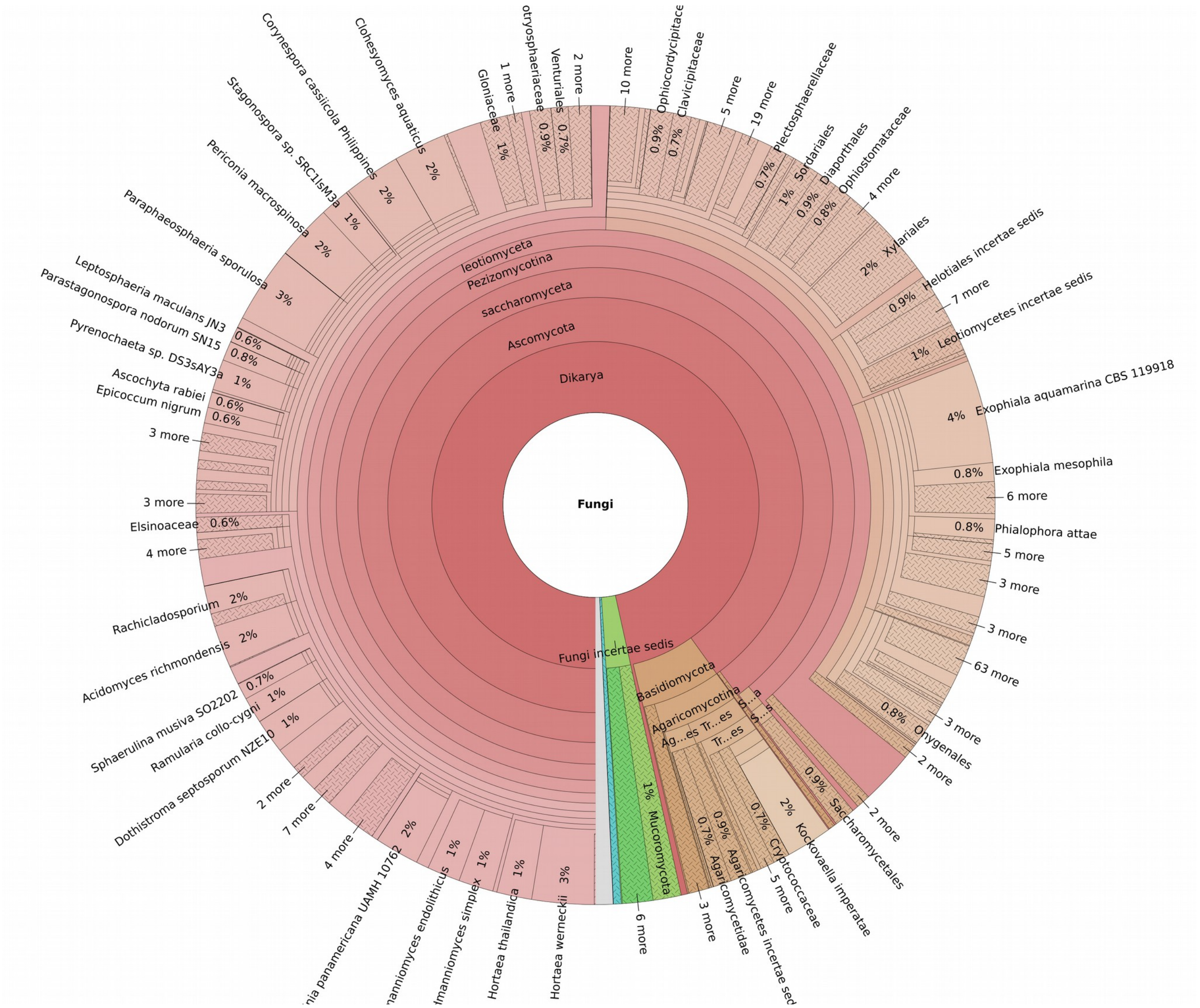


Figure S17. General overview of the fungal diversity in an avocado bark sample from Morelia (Sample M5).

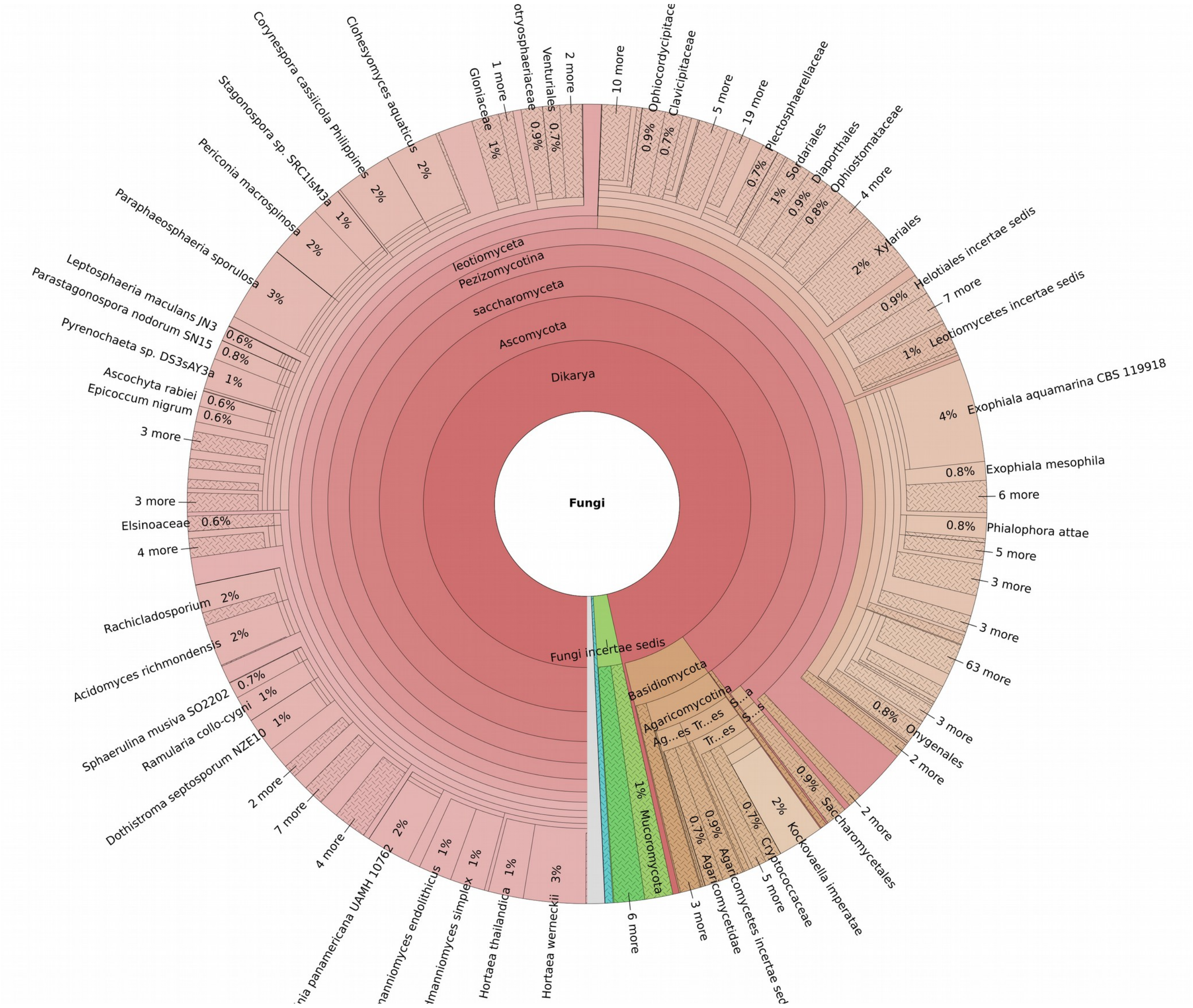

Figure S18. General overview of the fungal diversity in an avocado bark sample from Morelia (Sample M6).

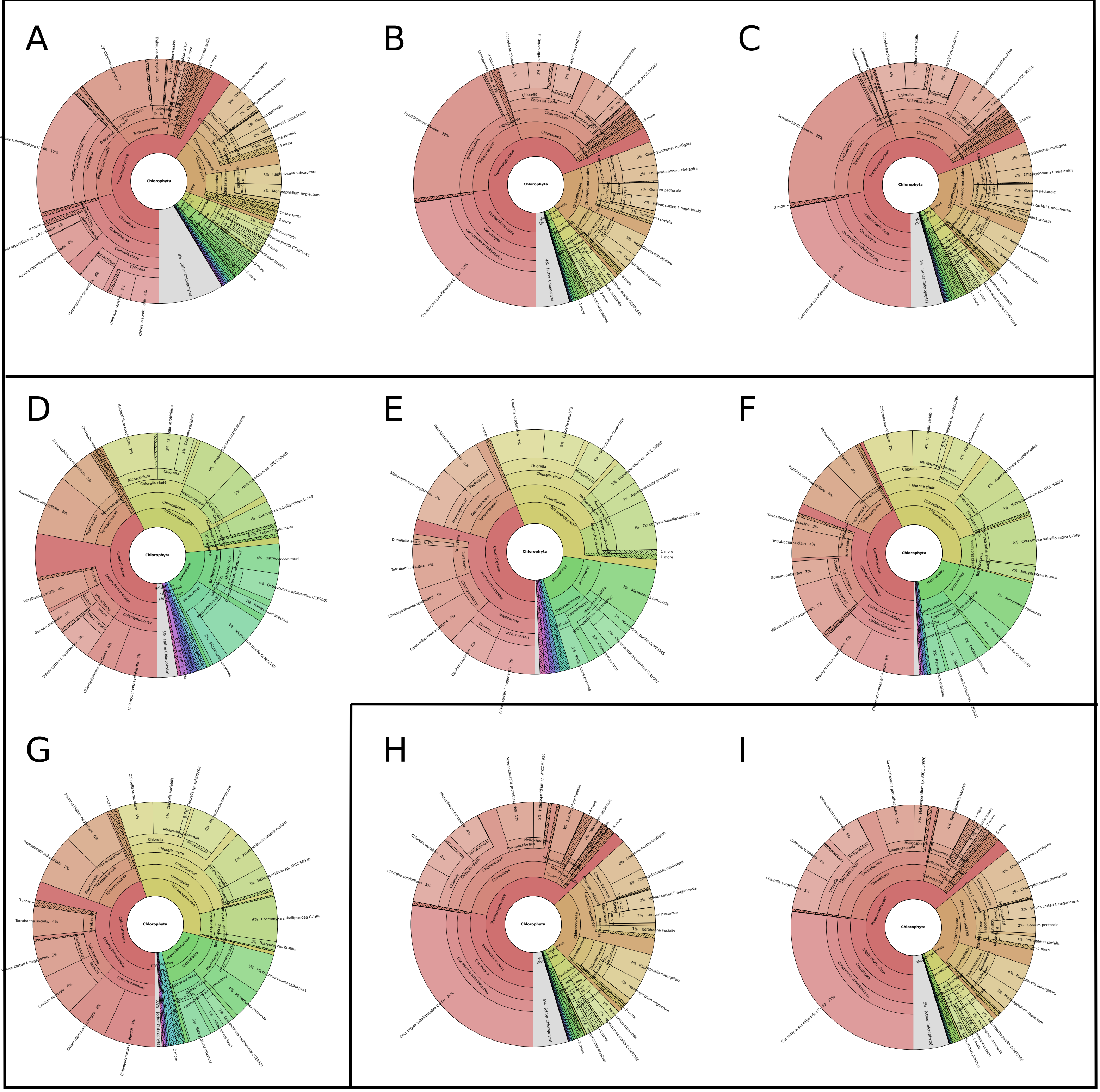


Figure S19. General overview of the diversity of Chlorophyta in an avocado bark and rhizosferic soil. A-C) Malinalco avocado bark, D-G) Malinalco rhizosferic soil, H-I) Morelia avocado bark.

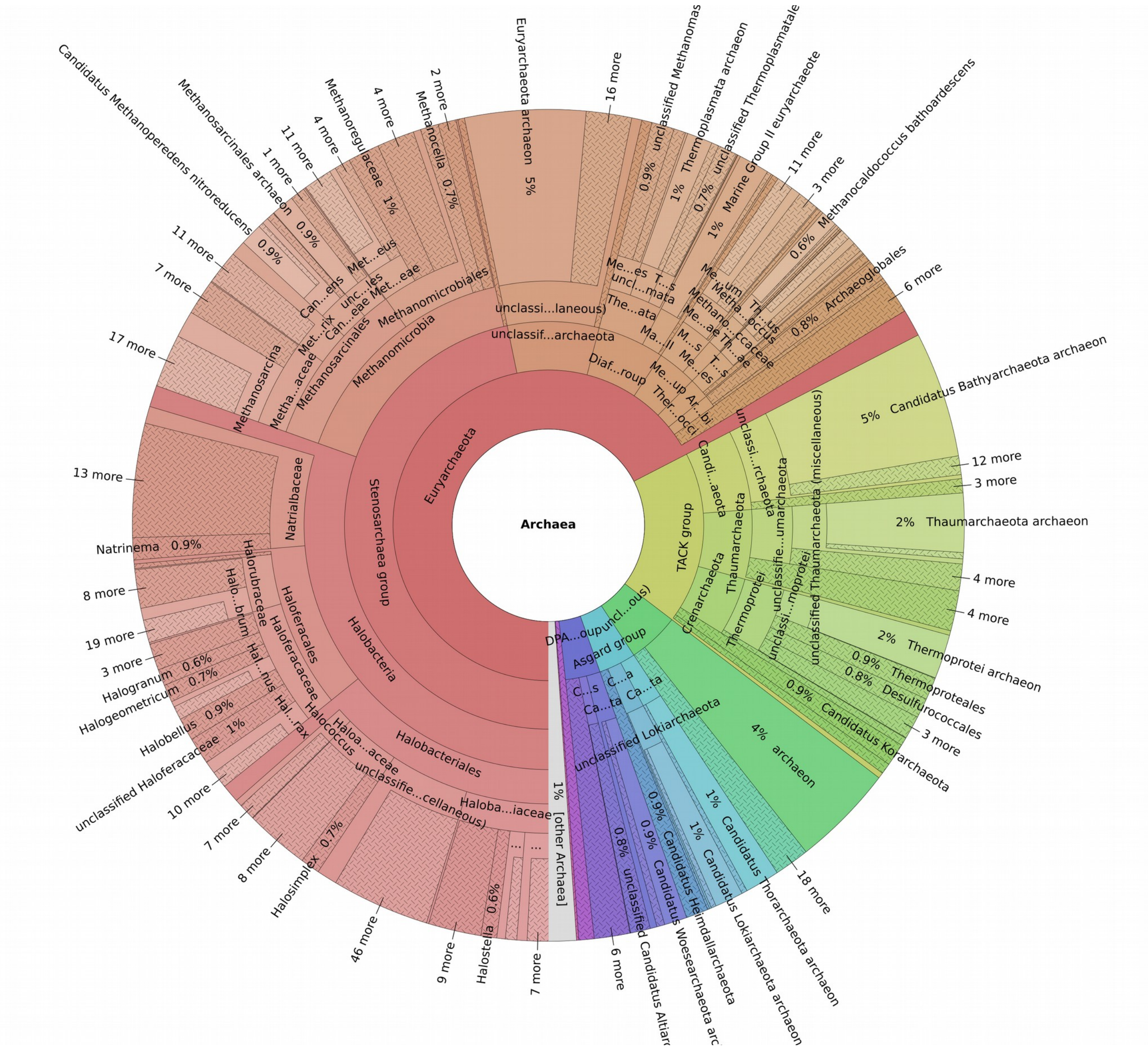

Figure S20. General overview of the archaeal diversity in an avocado bark sample from Malinalco (Sample A4).

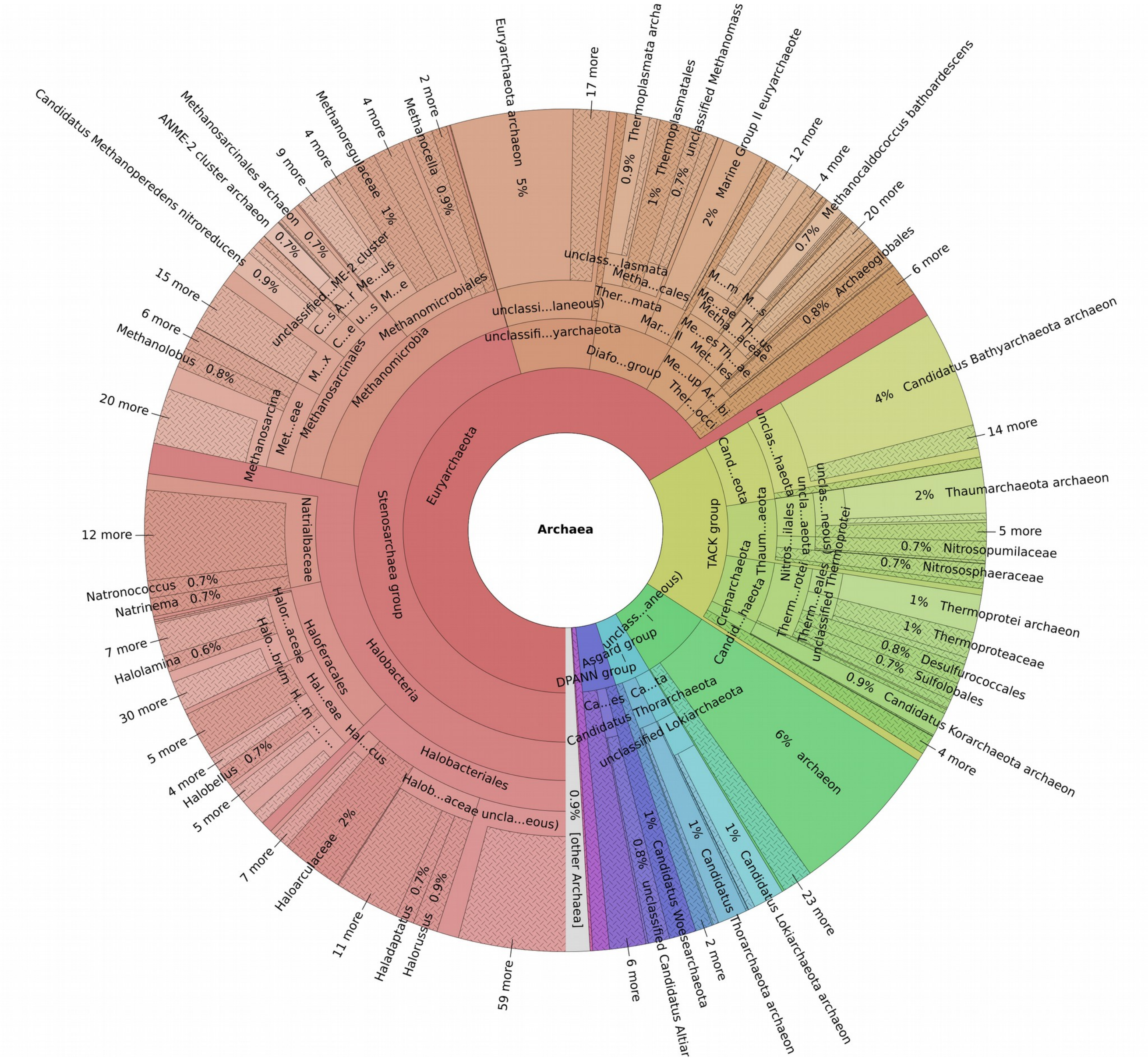

Figure S21. General overview of the archaeal diversity in an avocado bark sample from Malinalco (Sample A5).

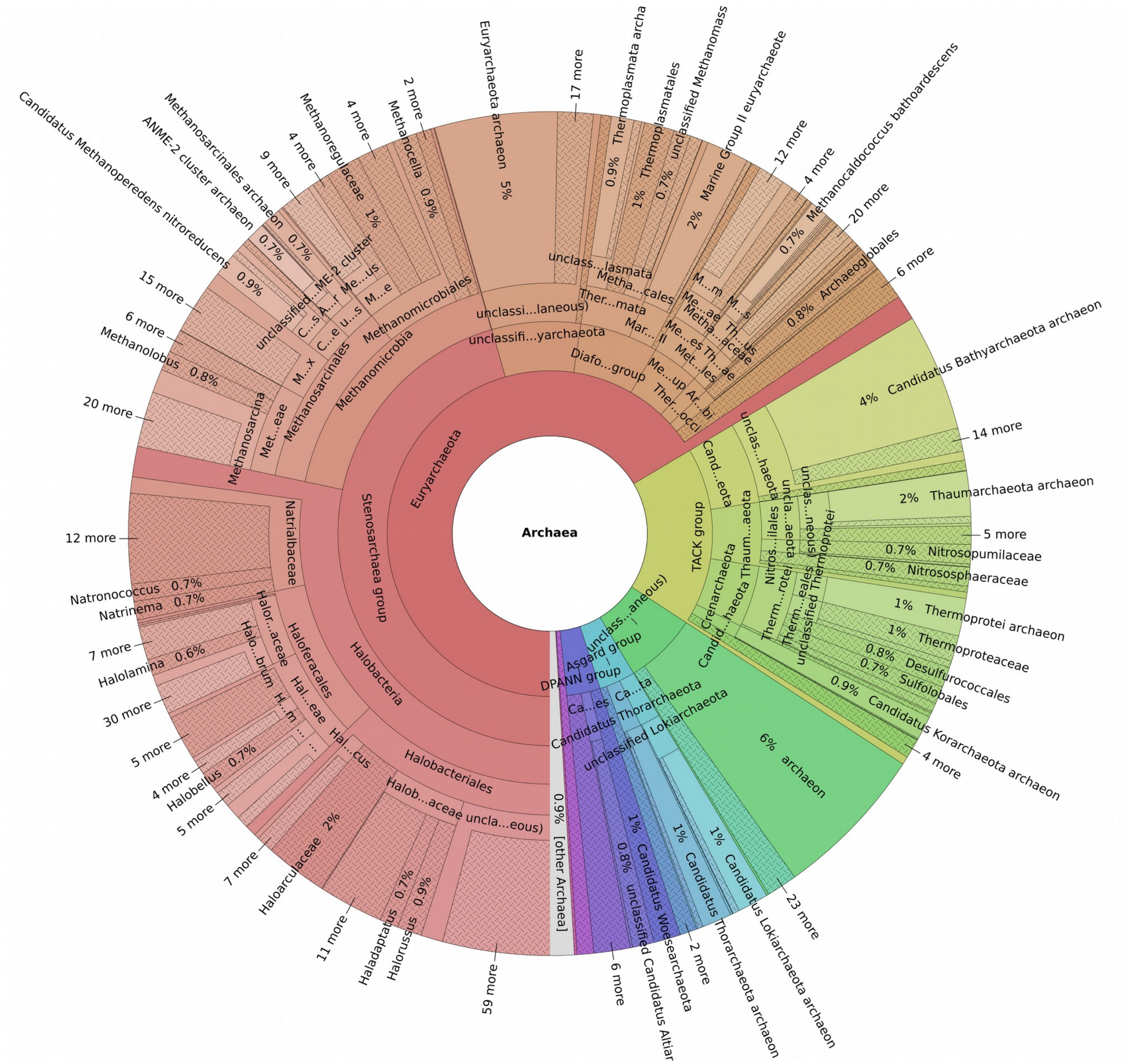

Figure S22. General overview of the archaeal diversity in a soil sample from Malinalco (Sample SA1).

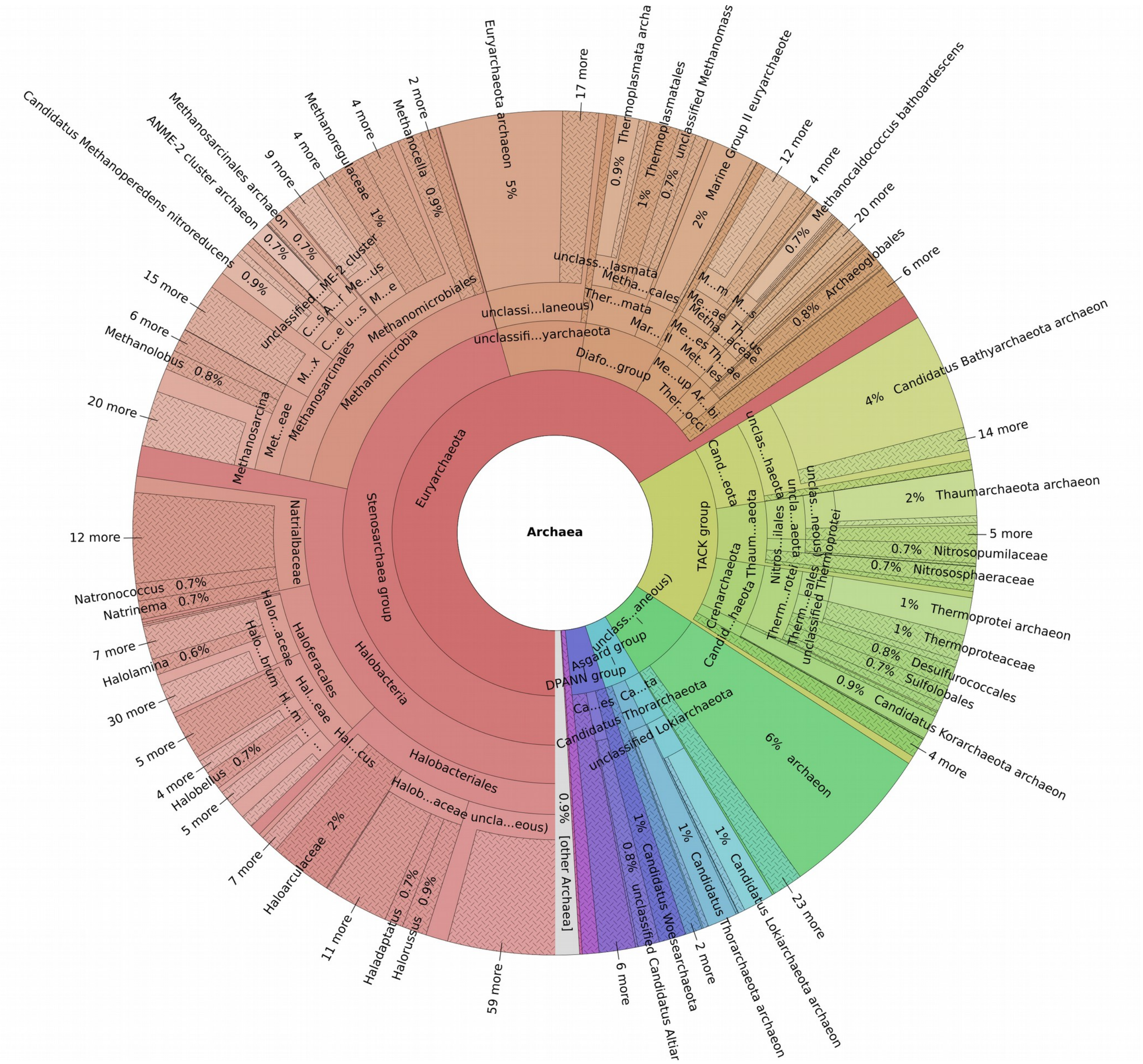

Figure S23. General overview of the archaeal diversity in a soil sample from Malinalco (Sample SA2).

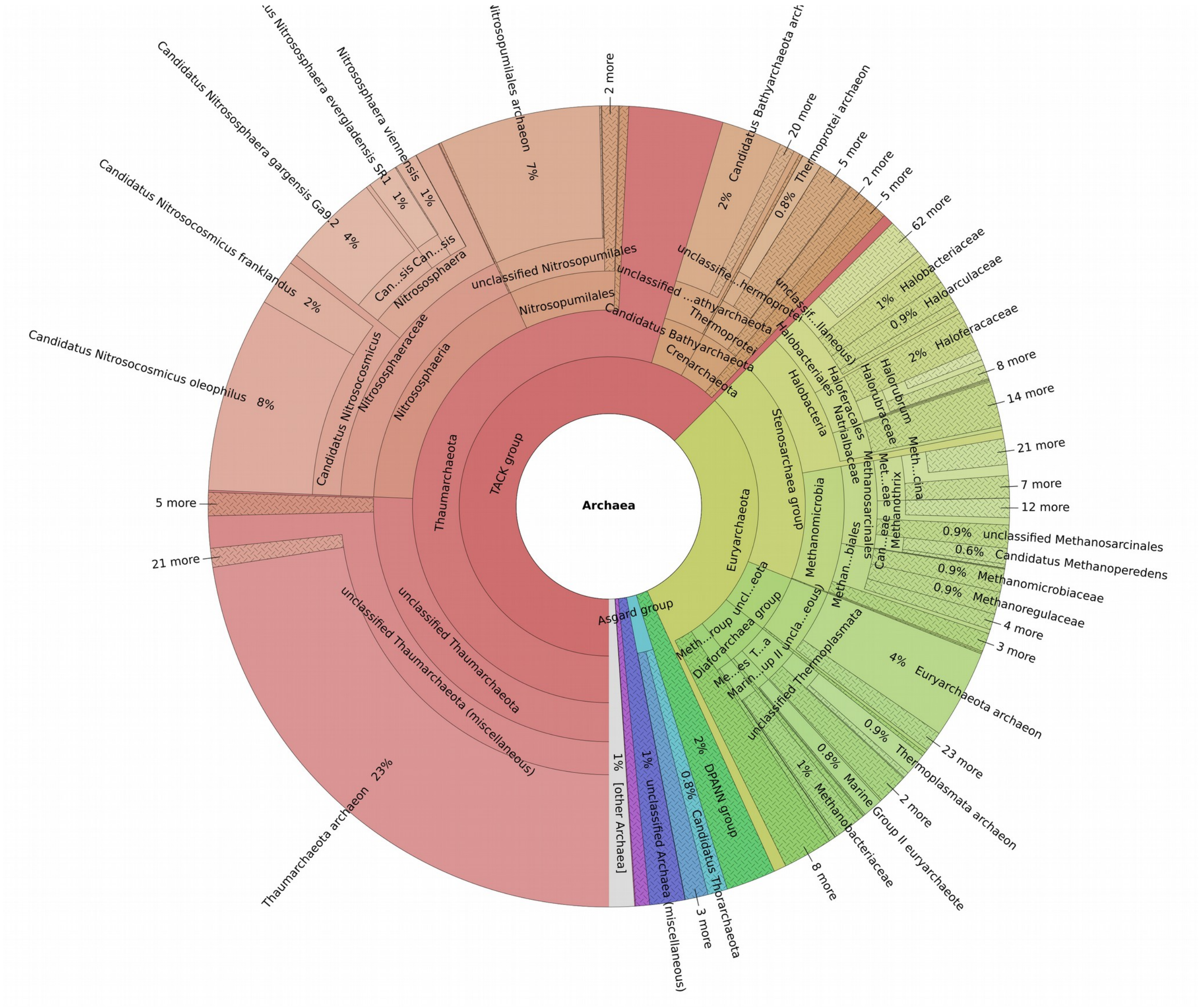

Figure S24. General overview of the archaeal diversity in a soil sample from Malinalco (Sample SA4).

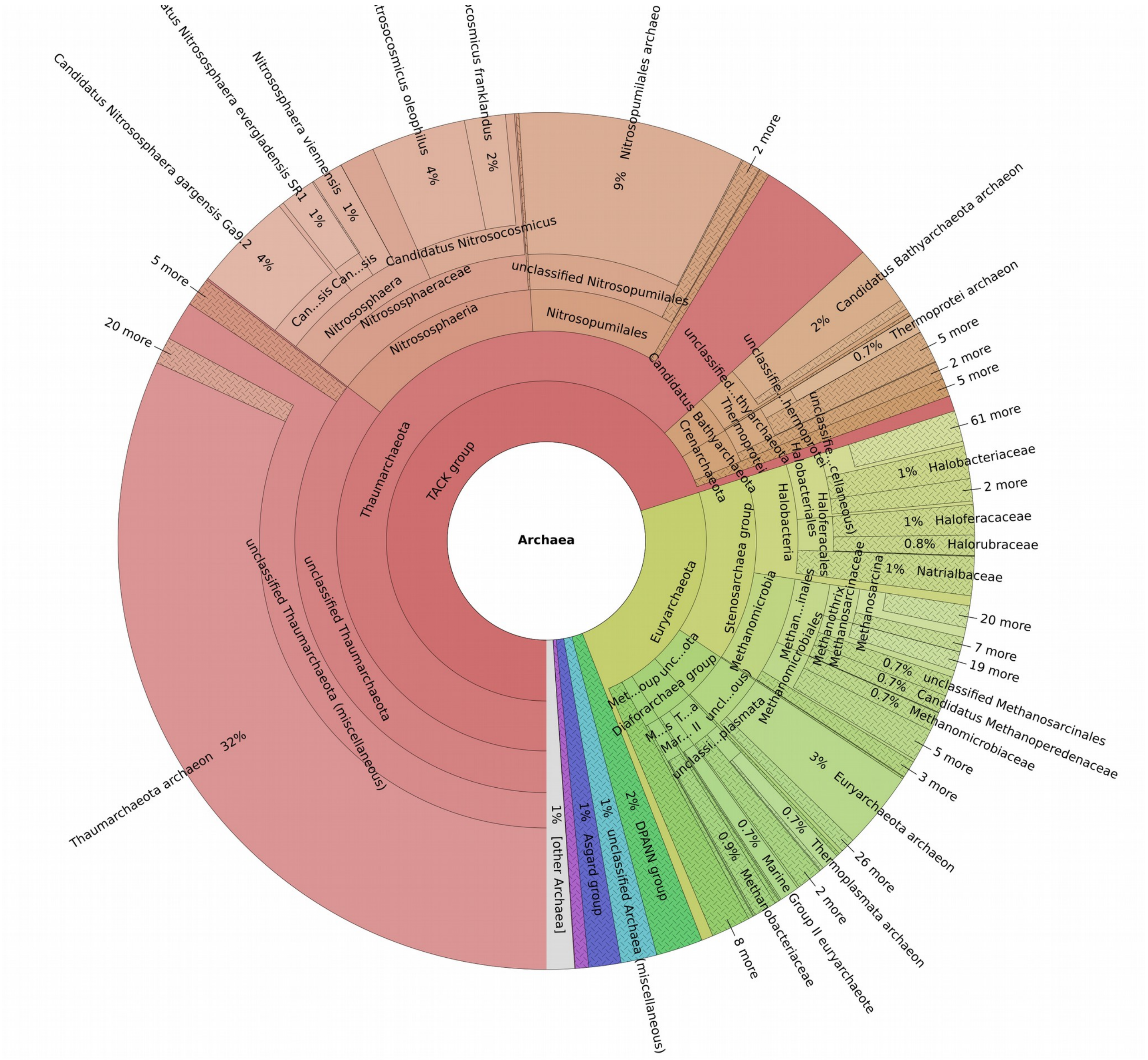


Figure S25. General overview of the archaeal diversity in a soil sample from Malinalco (Sample SA5).

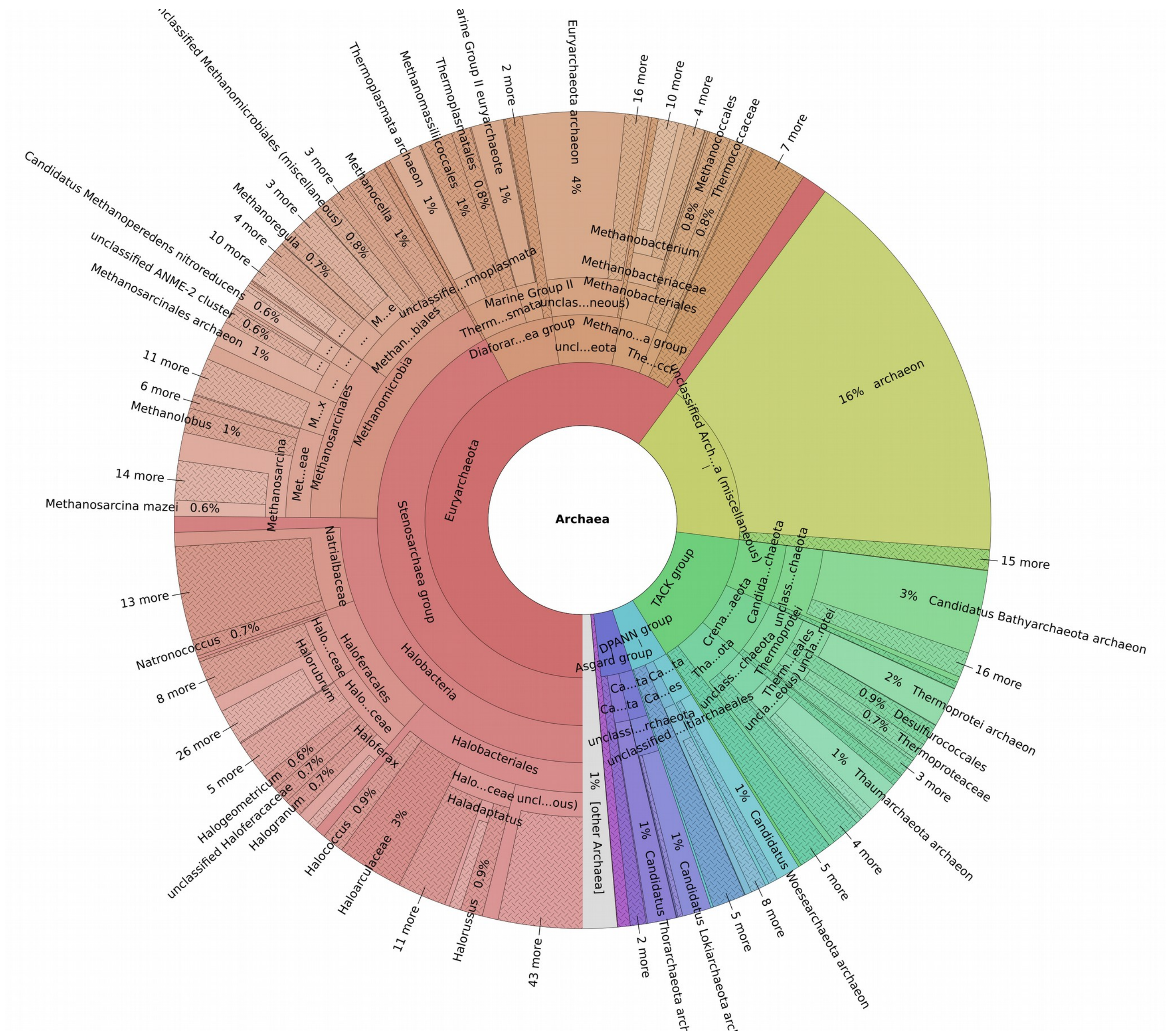

Figure S26. General overview of the archaeal diversity in an avocado bark sample from Morelia (Sample M5).

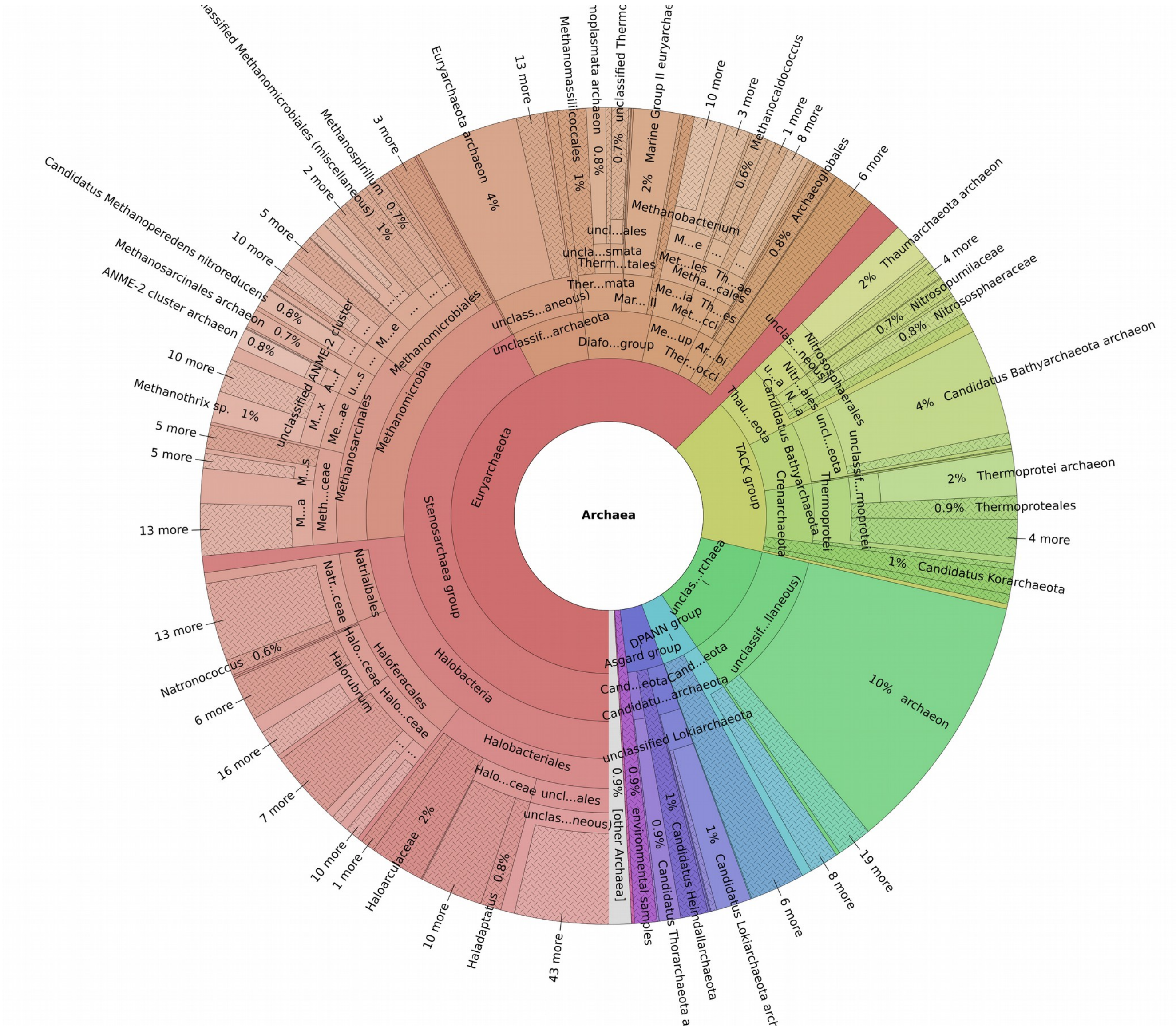


Figure S27. General overview of the archaeal diversity in an avocado bark sample from Morelia (Sample M6).

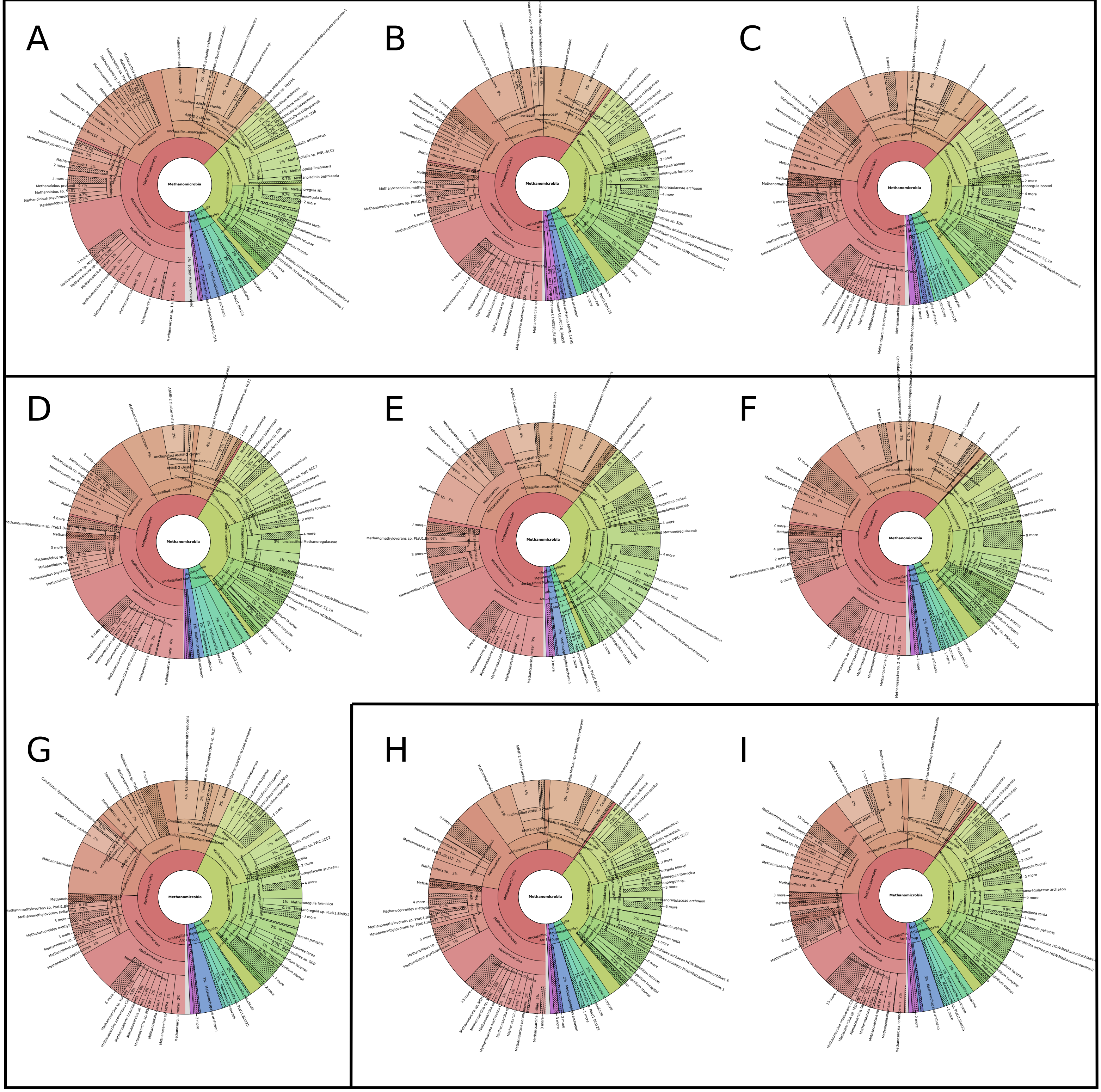


Figure S28. General overview of the diversity of the archaeal family Methanomicrobia in an avocado bark and rhizosferic soil. A-C) Malinalco avocado bark, D-G) Malinalco rhizosferic soil, H-I) Morelia avocado bark.

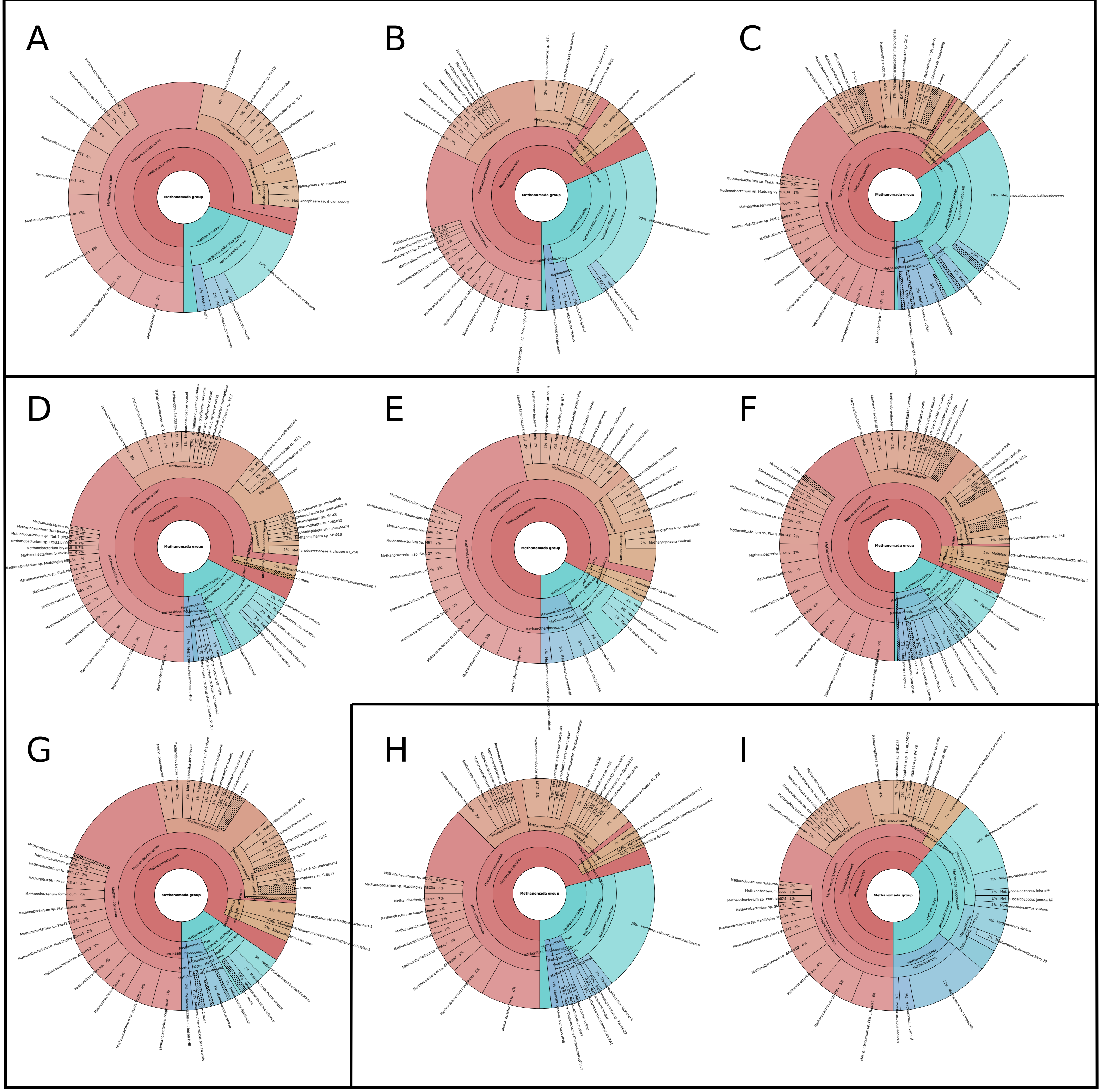


Figure S29. General overview of the diversity of the archaeal family Methanomada in an avocado bark and rhizosferic soil. A-C) Malinalco avocado bark, D-G) Malinalco rhizosferic soil, H-I) Morelia avocado bark.

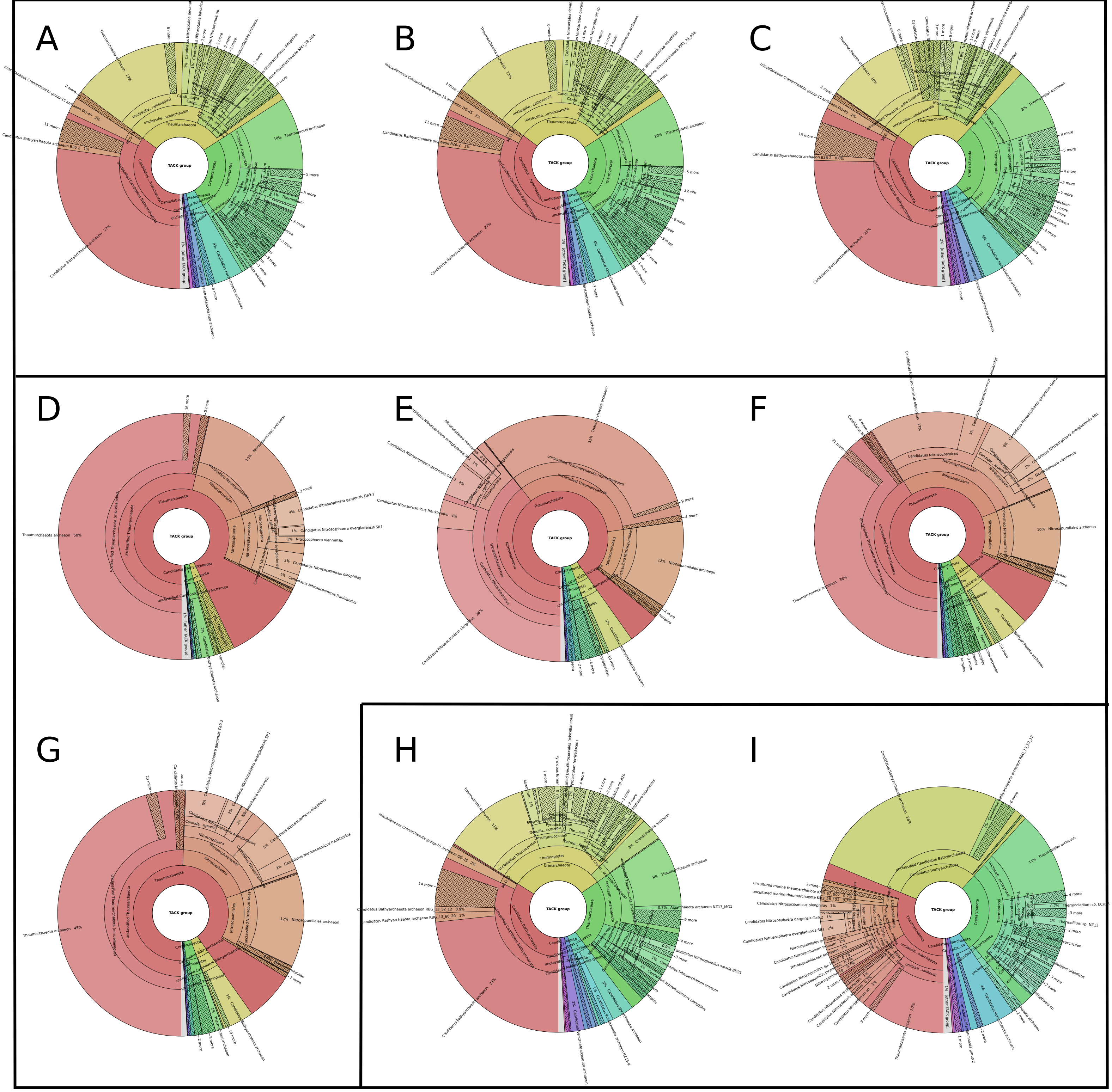

Figure S30. General overview of the diversity of the archaeal group TACK in avocado bark and rhizosferic soil. A-C) Malinalco avocado bark, D-G) Malinalco rhizosferic soil, H-I) Morelia avocado bark.
